# Pre-conditioning Modifies the Tumor Microenvironment to Enhance Solid Tumor CAR T Cell Efficacy and Endogenous Immunity

**DOI:** 10.1101/2020.11.18.388892

**Authors:** John P. Murad, Dileshni Tilakawardane, Anthony K. Park, Kelly T. Kennewick, Lupita S. Lopez, Hee Jun Lee, Brenna J. Gittins, Wen-Chung Chang, Chau P. Tran, Catalina Martinez, Anna M. Wu, Robert E. Reiter, Tanya B. Dorff, Stephen J. Forman, Saul J. Priceman

## Abstract

Chimeric antigen receptor (CAR) T cell therapy has led to impressive clinical responses in patients with hematological malignancies; however, its utility in patients with solid tumors has been limited. While CAR T cells for the treatment of advanced prostate cancer are being clinically evaluated and are anticipated to show bioactivity, their safety and the impact of the immunosuppressive tumor microenvironment (TME) have not been faithfully explored preclinically. Using a novel human prostate stem cell antigen knock-in (hPSCA-KI) immunocompetent mouse model and syngeneic murine PSCA CAR T cells, we performed analyses of normal and tumor tissues by flow cytometry, immunohistochemistry, and/or RNA sequencing. We further assessed the beneficial impact of cyclophosphamide (Cy) pre-conditioning on modifications to the immunosuppressive TME and impact on PSCA-CAR T cell safety and efficacy. We observed an *in vivo* requirement of Cy pre-conditioning in uncovering the efficacy of PSCA-CAR T cells in prostate and pancreas cancer models, with no observed toxicities in normal tissues with endogenous PSCA expression. This combination also dampened the immunosuppressive TME, generated pro-inflammatory myeloid and T cell signatures in tumors, and enhanced the recruitment of antigen-presenting cells, and endogenous as well as adoptively-transferred CAR T cells, resulting in long-term anti-tumor immunity.

## Introduction

Despite clinical successes of chimeric antigen receptor (CAR)-engineered T cell therapies in hematological malignancies, effective CAR T cell therapies in solid tumors has been limited.^1-3^ Immunotherapies for solid tumors are restricted by the tumor microenvironment (TME), which includes among others, tumor-associated macrophages (TAMs), myeloid-derived suppressor cells (MDSCs), and regulatory T cells (Tregs), all which suppresses endogenous immunity, as well as adoptively transferred T cell trafficking, persistence, and anti-tumor activity.^1,4,5^ Alleviating immunosuppression in solid tumors and improving CAR T cell-mediated anti-tumor activity is an active, but still very early, area of research and includes targeting immune checkpoint [e.g., PD-1 and CTLA-4] and/or other immunomodulatory pathways. Pre-conditioning chemotherapy has been widely employed in combination with CAR T cell therapy, particularly in the setting of hematological malignancies. This approach has classically been described as “space-making” lymphodepletion to enhance homeostatic cytokine production for improved adoptively-transferred T cell engraftment.^6,7^ However, the totality of benefits and underlying mechanisms of action of this approach for solid tumors is still controversial.^8^ Studies to more faithfully assess the safety and efficacy of this approach in the context of solid tumor CAR T cell therapies will require more comprehensive preclinical models.^9^

Our group has recently initiated a phase 1 clinical trial to evaluate the safety, feasibility and biological activity of prostate stem cell antigen (PSCA)-directed CAR T cells in patients with metastatic castration-resistant prostate cancer (mCRPC), based on extensive preclinical optimization using human xenograft models.^10^ As with most solid tumor-associated antigens (TAAs, e.g., HER2, CEA, PSMA, mesothelin), PSCA expression in normal tissues, including the stomach, bladder, pancreas, and prostate, may pose safety concerns or limit the therapeutic benefits of CAR T cells.^11,12^ While studies using immunocompromised mice allow for evaluating activity of CAR T cells *in vivo* for clinical translation, these immunocompromised mice typically lack a physiologically normal tissue expression of TAAs, which precludes assessment of potential “on-target, off-tumor” toxicities. Additionally, they fail to capture the complexity of the local TME and the impact of immunotherapy on systemic immunity. These components may contribute to some of the key discrepancies in clinical CAR T cell responses observed in hematological malignancies and solid tumors.^13,14^

In the current study, we developed an immunocompetent mouse model that allows simultaneous assessment of safety and anti-tumor efficacy of PSCA-CAR T cells. Our knock-in mouse system allowed for expression of human PSCA under the control of the mouse PSCA promoter in normal tissues. This model enabled us to evaluate PSCA-CAR T cell therapy in the context of a host with an intact immune system and the ability to comprehensively interrogate mechanisms underlying response and/or resistance to CAR T cell therapy. We found PSCA-CAR T cells ineffective in eliciting *in vivo* anti-tumor responses unless given after lymphodepleting pre-conditioning with cyclophosphamide (Cy). Mechanistically, we showed that the benefits of Cy pre-conditioning were attributed to early changes in the TME, including pro-inflammatory myeloid cell modifications, improved antigen presentation pathways, and profound tumor infiltration of both endogenous and adoptively-transferred T cells. Combining Cy pre-conditioning with PSCA-CAR T cells resulted in durable curative responses and subsequent protective immunity against tumor rechallenge. Importantly, potent PSCA-CAR T cell anti-tumor responses were not associated with adverse effects on normal tissues expressing PSCA, or other overt off-tumor toxicities. These findings will inform our ongoing clinical trials and further provide a preclinical platform to design rational combination approaches to maximize the safety and efficacy of immunotherapies for PSCA+ solid tumors.

## Results

### Murine PSCA-CAR T cells demonstrate selective *in vitro* activation against PSCA+ murine tumor cells

We generated a fully-murine PSCA-CAR retroviral construct (PSCA-mCAR) targeting human PSCA. Importantly, the human PSCA targeting scFv used (clone 1G8) is the same clone that was subsequently humanized to generate our current phase 1 clinical trial (NCT03873805) lead therapeutic candidate (**Figure 1 a**).^10^ PSCA-mCAR retrovirus yielded efficient transduction of murine splenic CD4/CD8 T cells as determined by expression of mCD19t (**Figure 1 b**). *In vitro* antigen-specific activity of PSCA-mCAR T cells was assessed by co-culture of wild-type RM9 (hPSCA-) or RM9-hPSCA tumor cells with freshly-transduced PSCA-mCAR T cells, resulting in antigen-dependent secretion of murine IFNγ and IL-2 cytokines (**Figure 1 c and d**). PSCA-mCAR T cells also showed antigen-dependent activation and exhaustion markers murine 4-1BB and PD-1, respectively, at varying effector:tumor (E:T) ratios (**Figure 1 e**). RM9-hPSCA tumor cells exhibited CAR T cell-dependent increases in PD-L1 *in vitro*, presumably in response to IFNγ secretion by activated CAR T cells (**Figure 1 f**). CAR T cell-mediated killing of RM9-hPSCA tumor cells increased from 24 h and 72 h (**Figure 1 g**). Additionally, PSCA-mCAR T cells also expanded following 72 h co-culture (**Figure 1 h**). These data show that our fully-murine PSCA-CAR T cells exhibit strong antigen-specific activity against murine tumors expressing human PSCA.

**Figure 1:**
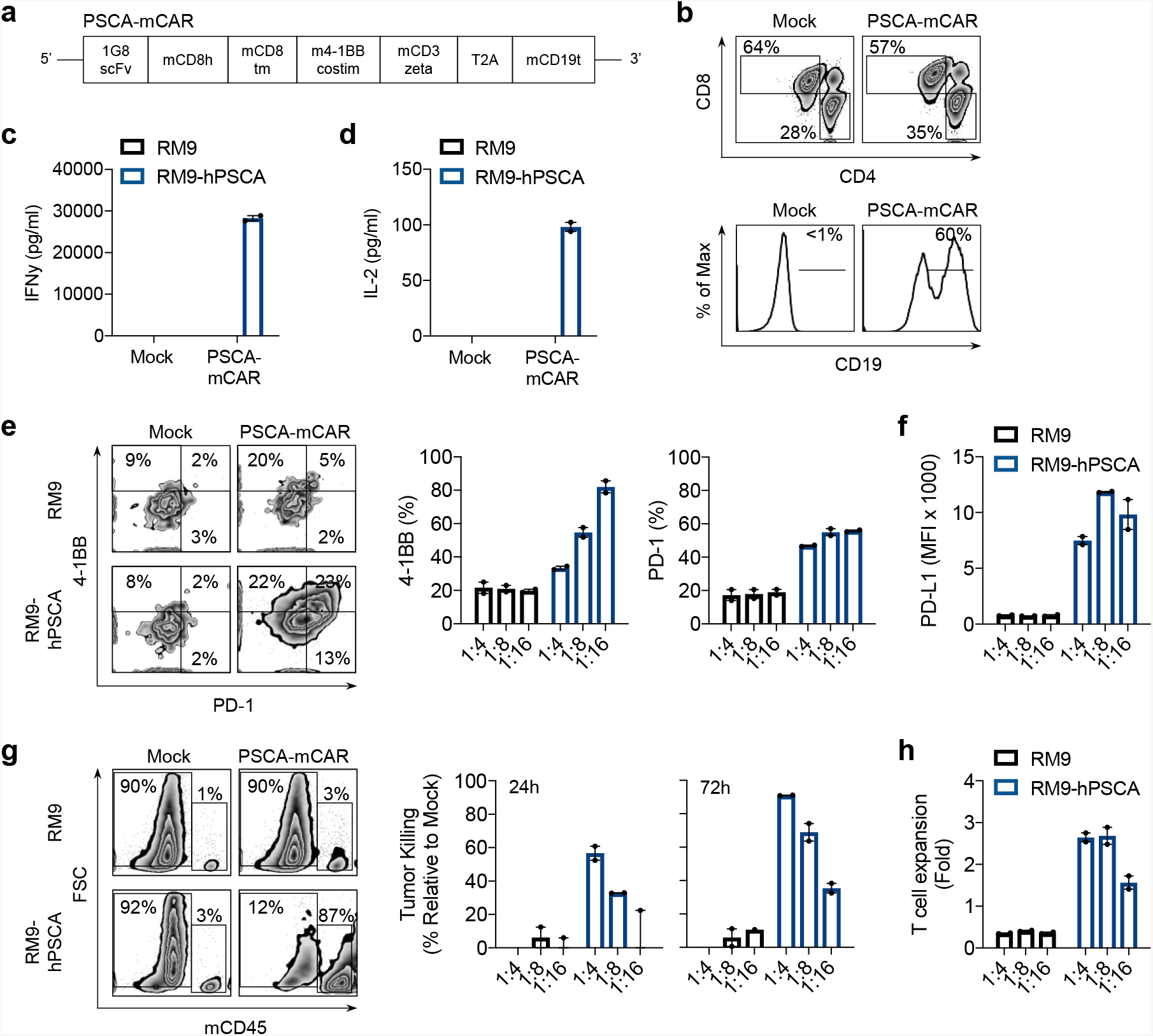
Characterization of PSCA-mCAR T cell transduction and activity *in vitro*. a) Diagram of the fully murine retroviral expression cassette with PSCA-mCAR T cell containing murine scFv (1G8 clone) targeting human PSCA. A truncated non-signaling murine CD19 (CD19t), separated from the CAR sequence, was expressed for identifying transduced T cells. b) Flow cytometry detection of CAR transduction measured by CD19t expression on Mock (untransduced) and PSCA-mCAR T cells (top) and CD4 and CD8 expression (bottom). IFNγ (c) and IL-2 (d) secretion, as measured by ELISA, by Mock and PSCA-mCAR T cells after 24 h co-culture with antigen-negative RM9 and antigen-positive RM9-hPSCA. For ELISA, N≥2 replicates per group. e) Representative flow cytometry plots depicting %4-1BB and %PD-1 induction on Mock or PSCA-mCAR after 72 h co-culture with RM9 or RM9-hPSCA (left) and quantification of %4-1BB and %PD-1 expression at indicated E:T ratios (right). f) Quantification of mean fluorescent intensity (MFI) of PD-L1 expression on indicated cancer cells after 72 h co-culture with Mock or PSCA-mCAR T cells. g) Representative 72 h flow cytometry plots of co-cultures of RM9 or RM9-hPSCA with Mock or PSCA-mCAR T cells showing dapi negative, CD45-negative (remaining viable tumor cells) or CD45-positive (T cell expansion) (left), and quantification of killing at 24 and 72 h post co-culture at indicated E:T ratios (right). h) Quantification of fold expansion of Mock or PSCA-mCAR T cells after 72 h co-culture with RM9 or RM9-hPSCA. For co-culture flow cytometry data, N≥3 replicates per group. All data are representative of two independent experiments.

### PSCA-mCAR T cells lack *in vivo* therapeutic efficacy in immunocompetent mice

To investigate the safety and efficacy of PSCA-CAR T cells in an immunocompetent system, we utilized a recently developed human PSCA knock-in (hPSCA-KI) mouse model.^15^ We used heterozygous mice (hPSCA-KI^het^) (**Figure S1 a**), to allow for engraftment of RM9-hPSCA tumor cells that may express both endogenous murine and engineered human PSCA. Using flow cytometry, immunohistochemistry (IHC), and (RNAscope™) analyses of normal and tumor tissue, we observed low to moderate levels expression of hPSCA, relative to RM9-hPSCA tumors, in normal prostate epithelia, bladder, and stomach, mimicking expression patterns observed in humans (**Figure S1 b-d**).^15,16^

Safety and efficacy of PSCA-CAR T cells were assessed in hPSCA-KI mice bearing s.c. RM9-hPSCA tumors treated with either Mock (untransduced) or PSCA-mCAR T cells by i.v. delivery. In contrast to the potent activity seen in previously published human xenograft models,^10^ we observed minimal anti-tumor responses in hPSCA-KI mice treated with PSCA-mCAR T cells (**Figure S2 a**). Encouragingly, no overt toxicities including visual weight loss or gross anatomical defects to normal organs upon euthanasia were observed in these mice. Interestingly, lymphocyte-deficient RAG2^-/-^ mice bearing RM9-hPSCA tumors treated with singularly with PSCA-mCAR T cells demonstrated curative responses in 33% of treated mice (**Figure S2 b**), highlighting the therapeutic potential of these CAR T cells in immunocompromised mice, and also suggesting that in immunocompetent mice, the intact endogenous immune system is likely contributing to a suppressive TME which resulted in limited efficacy of PSCA-mCAR T cells alone.

### *In vivo* efficacy of PSCA-mCAR T cells in immunocompetent mice requires cyclophosphamide preconditioning

Given the observed efficacy of PSCA-mCAR T cells in lymphocyte-deficient RAG2^-/-^ mice but not in immunocompetent hPSCA-KI mice (**Figure S2 a and b**), we assessed whether lymphodepleting pre-conditioning could improve the therapeutic impact of PSCA-mCAR T cells in this model. An *in vivo* Cy dose titration using a single intraperitoneal (i.p.) dose of Cy (50, 100, or 200 mg/kg) in hPSCA-KI mice resulted in a dose-dependent peripheral lymphodepletion in each immune subset evaluated, with a maximal lymphodepletion achieved 4 days following Cy treatment (**Figure S3 a - e**). The anti-tumor activity of Cy alone was then examined in i.ti.tumor bearing hPSCA-KI mice. From these studies we determined that in this model a single i.p. 100 mg/kg dose of Cy could provide adequate peripheral lymphodepletion, with minimal anti-tumor activity, and an absence of survival benefit (**Figure S3 f**).

We next evaluated the impact of 100 mg/kg Cy pre-conditioning in combination with PSCA-mCAR T cell therapy *in vivo*. PSCA-mCAR T cells or Cy treatment alone showed essentially no or only transient anti-tumor activity against RM9-hPSCA tumors respectively, and neither provided durable survival benefits (**Figure 2 a**). However, the combination of PSCA-mCAR T cells with Cy pre-conditioning elicited robust anti-tumor activity, improved overall survival, and complete responses (CR) in over 40% of mice (**Figure 2 b and c**). These data strongly suggest that pre-conditioning in this model is required to unleash the therapeutic potential of PSCA-mCAR T cells. Importantly, no overt toxicities or gross changes in cellular architecture to hPSCA expressing normal tissues (prostate, bladder, stomach) were observed at early and later timepoints post T cell treatment in any of the treated mice, including the combination of PSCA-mCAR T cells and Cy (**Figure 2 d**).

**Figure 2:**
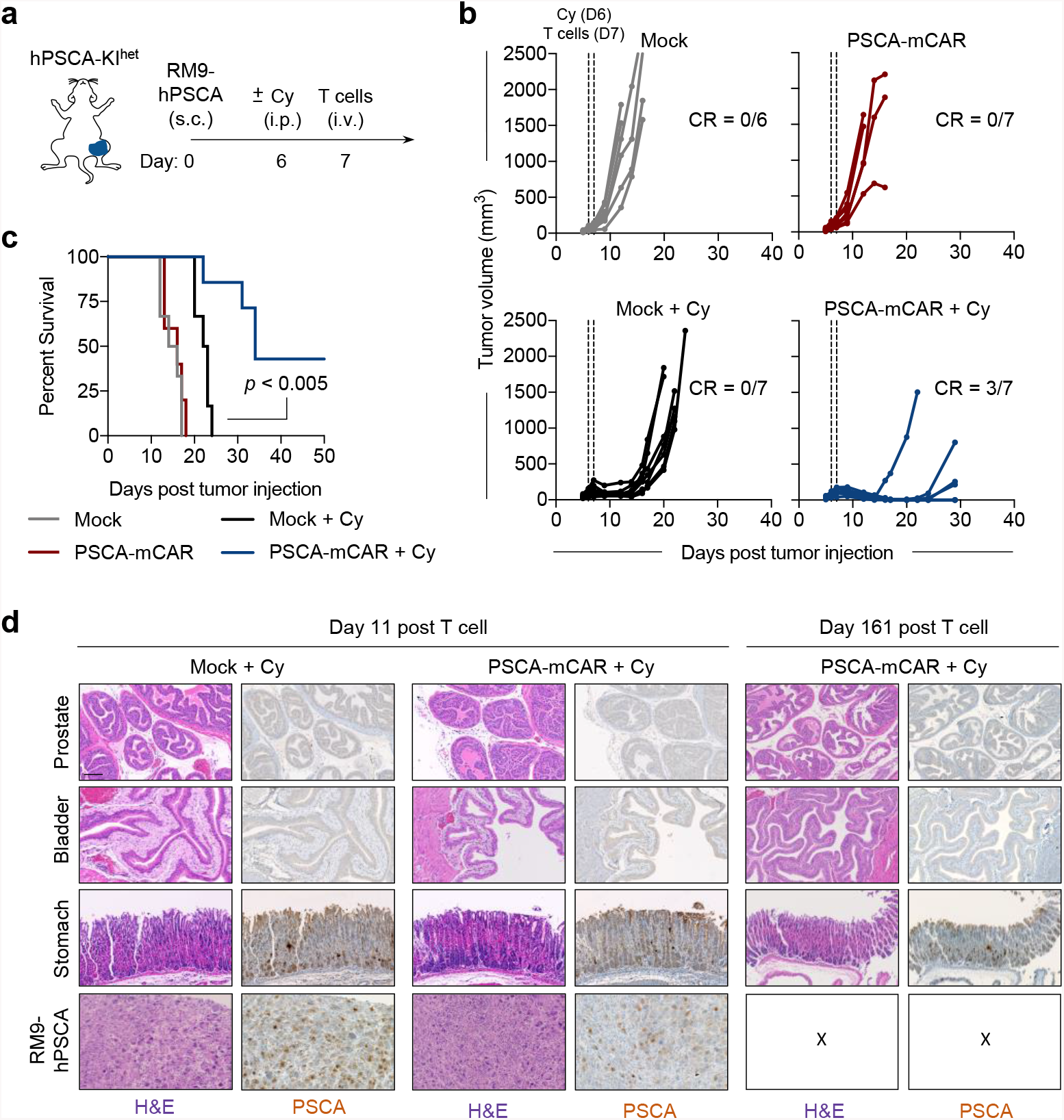
Anti-tumor effect of PSCA-mCAR following Cy mediated pre-conditioning and histological evaluation of normal tissue architecture and PSCA expression following treatment *in vivo*. a) Illustration of RM9-hPSCA s.c. tumor engraftment and Cy pre-conditioning (vehicle or Cy 100mg/kg, i.p.) 24 h prior to treatment with 5.0×10^6^ T cells (Mock or PSCA-mCAR, i.v.). b) Tumor volume (mm^3^) measurements of each replicate at indicated days post tumor injection for indicated treatment with or without Cy pre-conditioning; *CR = complete response*. c) Kaplan-Meier survival plot for mice in each group indicated. N≥6 mice per group. d) Representative IHC (H&E and PSCA protein staining) of indicated normal tissues from Cy-preconditioned tumor bearing mice at day 11 (left) or day 161 (right) post Mock or PSCA-mCAR T cells injection. All images at 20x magnification, scale bar represents 100 μm.

### Cyclophosphamide reverts T cell exclusion and promotes tumor infiltration of CAR T cells

We next evaluated the effects of Cy on the local TME. Tumors were initially harvested at day 3 post CAR T cell therapy and analyzed by IHC. Interestingly, a clear T cell exclusion phenotype was observed in tumors from mice treated with Mock or PSCA-mCAR T cells alone, with T cells relegated to the normal tissue periphery and only few T cells within the tumor (**Figure 3 a**). Following Cy pre-conditioning, T cells were observed at a much higher frequency within the tumor, which was greatly enhanced in combination with PSCA-mCAR T cell treatment. These data, coupled with impressive *in vivo* anti-tumor responses with the combination (**Figure 2**), suggest that Cy may revert T cell exclusion within the TME.

**Figure 3:**
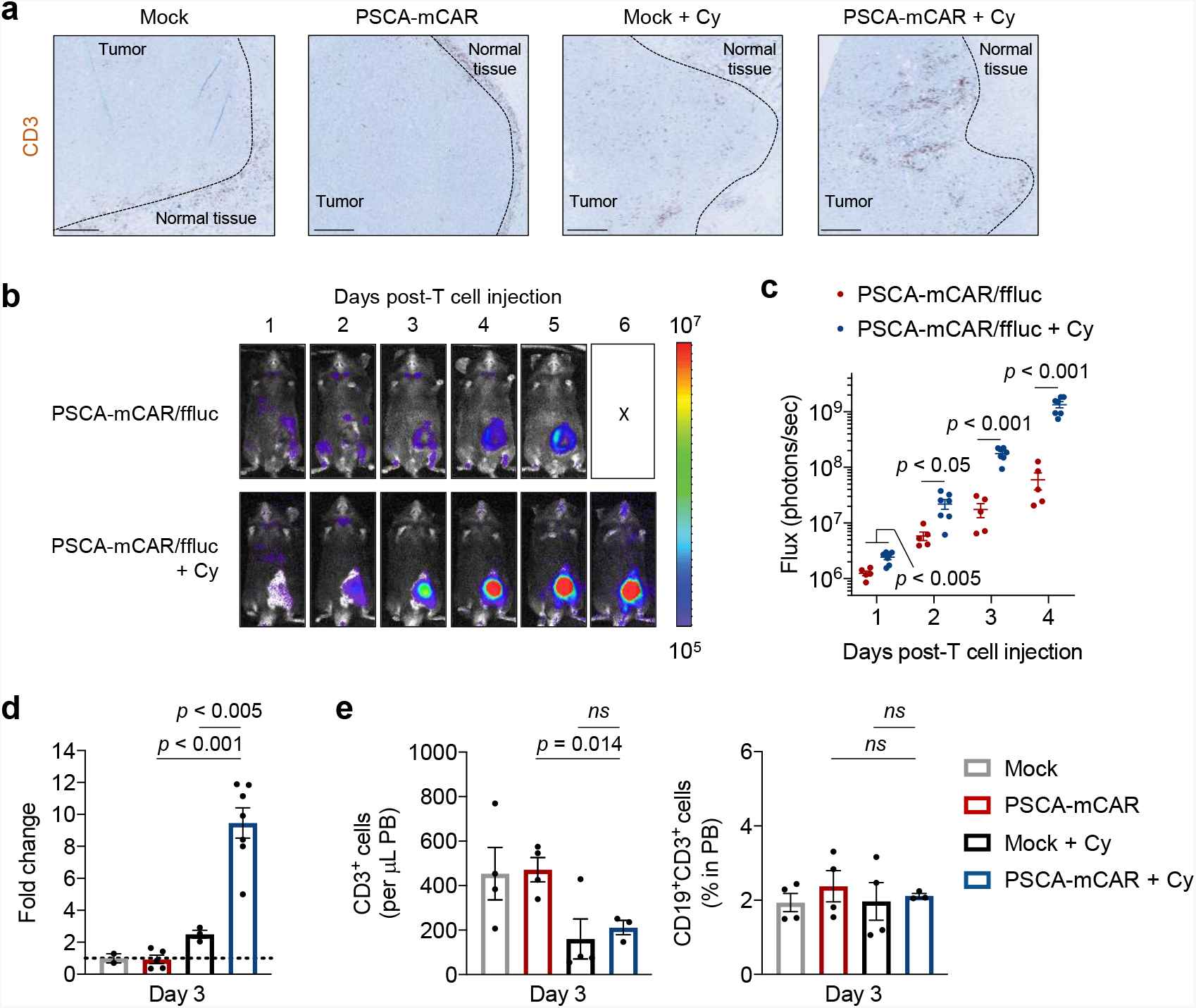
Cy pre-conditioning improves intra-tumoral PSCA-mCAR T cell accumulation and expansion *in vivo*. a) IHC of CD3 T cell localization in representative RM9-hPSCA tumors harvested 3 days post T cell injection in all treatment groups. b) Time-course bioluminescent flux imaging in RM9-hPSCA tumor bearing mice following injection of *ffluc*-positive PSCA-mCAR T cells with or without Cy pre-conditioning. c) Quantification of PSCA-mCAR/ffluc T cell flux on days 1 through 4 post T cell injection with or without Cy pre-conditioning. N≥5 mice per group. d) Fold change in PSCA-mCAR/ffluc T cell flux on day 3 post T cell injection from panel b and c. e) Flow cytometry quantification of total CD3+ T cell counts per μL PB (left) and %PSCA-mCAR (%CD19^+^ gated on total CD3^+^) found in PB (right) collected from mice on day 3 post T cell injection with or without Cy pre-conditioning. N≥5 mice per group. IHC 5x magnification, scale bar represents 500 μm. All data are representative of two independent experiments.

We then quantified the kinetics of PSCA-mCAR T cell trafficking with or without Cy pre-conditioning. hPSCA-KI mice s.c. engrafted with RM9-hPSCA (non-firefly luciferase expressing) received Cy pre-conditioning followed 24 h later with i.v. firefly luciferase-expressing Mock/ffluc or PSCA-mCAR/ffluc T cells. Mice were imaged daily to monitor CAR T cell biodistribution (**Figure 3 b**). We observed significant increases of intratumoral PSCA-mCAR T cell flux in Cy pre-conditioned mice as early as 24 h post T cell treatment (**Figure 3 c**) as compared with mice treated with PSCA-mCAR T cells alone where T cell signal was predominantly confined at the tumor periphery. Interestingly, the early intratumural accumulation of CAR T cells was observed prior to maximal Cy-mediated peripheral lymphodepletion in our model. By day 3 post CAR T cell treatment, intratumoral proliferation/expansion of CAR T cells following Cy pre-conditioning was nearly 10-fold higher than with CAR T cells alone (**Figure 3 d**). Furthermore, analysis of PB at days 1, 3 or 10 post T cell treatment, and tumor-draining lymph nodes at 7 days post T cell treatment, both failed to show an accumulation of peripheral PSCA-mCAR T cells, despite evidence of a Cy mediated peripheral lymphodepletion (**Figure 3 e, Figure S4 a and b**). These data highlight a Cy-mediated modulation of the local TME resulting in tumor-specific trafficking and expansion of PSCA-mCAR T cells.

### Cyclophosphamide overcomes the immunosuppressive TME

To better understand the impact of Cy on the local TME, we performed RNA sequencing of tumors in each treatment group. Analysis of standardized transcript expression revealed that Cy pre-conditioning induced widespread transcriptional changes in comparison to Mock or PSCA-mCAR T cells alone, indicating a local tumor modulating effect of Cy (**Figure 4 a**). A gene ontology (GO) Gene Set Enrichment Analysis (GSEA) of biological processes was performed and identified most impacted gene pathways. GSEA analysis showed significant enrichment of T cell migration pathways in tumors following Cy pre-conditioning alone (**Figure 4 b, top**). Tumors from PSCA-mCAR T cells in combination with Cy showed multiple pathway enrichment sets, highlighted by multiple T cell activation pathways, increased IFNγ production, increased adaptive immune responses, and monocyte chemotaxis (**Figure 4 b, bottom**).

**Figure 4:**
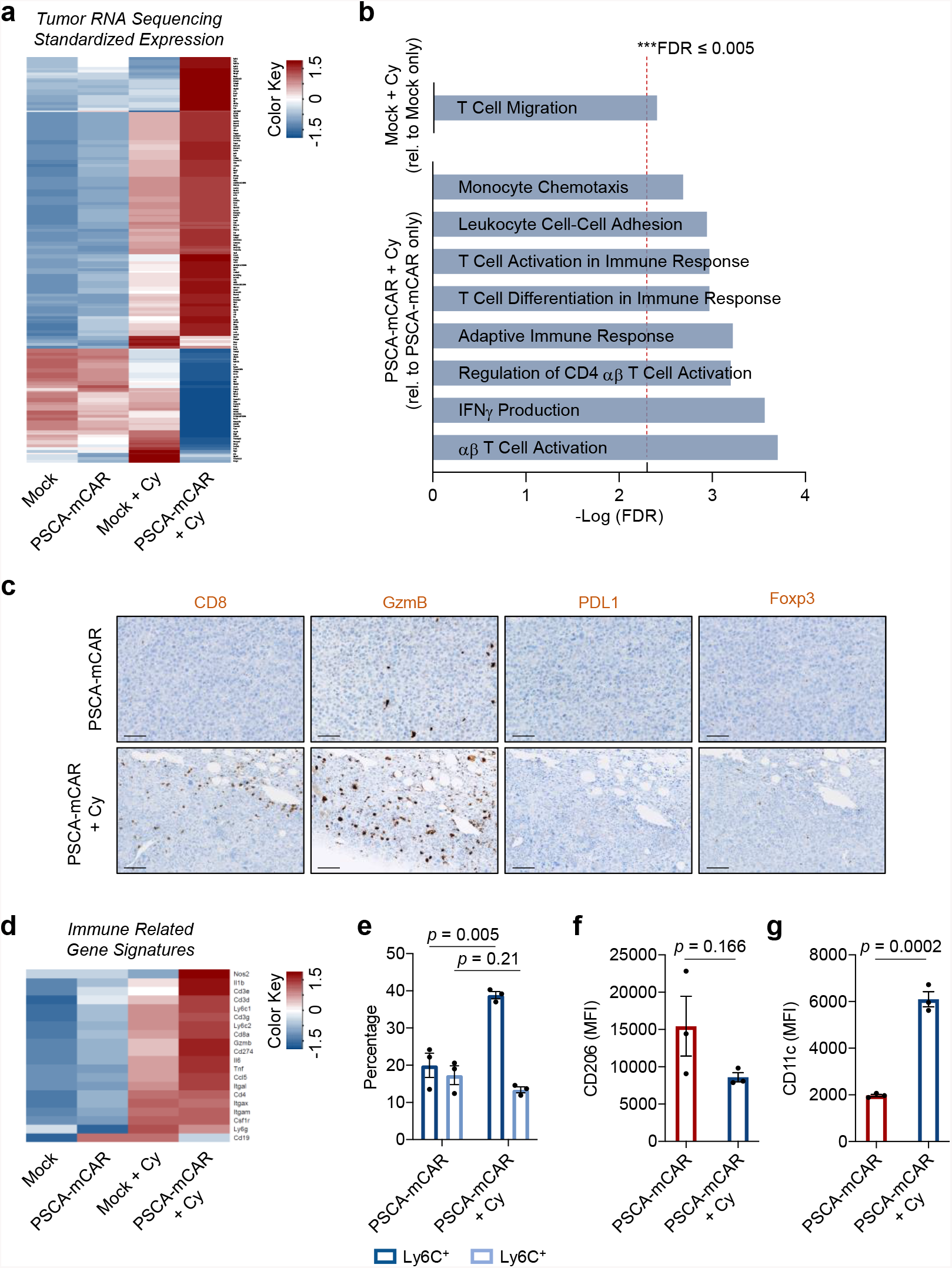
Tumor RNA sequencing analysis reveals Cy mediated pro-inflammatory modulation of the TME. a) RNA sequencing heatmap showing standardized expression of transcripts from RM9-hPSCA tumors after Mock or PSCA-mCAR treatment with or without Cy pre-conditioning (scale −1.5 to +1.5). b) GSEA network GO pathway analysis highlighting significant enrichment (FDR ≤ 0.005) in immune related biological processes from tumors treated with Mock + Cy relative to Mock alone (top) or PSCA-mCAR + Cy treatment relative to PSCA-mCAR alone (bottom). c) IHC of CD8, GzmB, PDL1, and Foxp3 in representative tumor tissues treated with PSCA-mCAR T cells alone or PSCA-mCAR following Cy pre-conditioning. 20x magnification, scale bars represent 100 μm. d) Heatmap showing standardized expression of immune related gene signatures from bulk tumors reveal pro-immune shifts following PSCA-mCAR or combo treatment (scale −1.5 to +1.5). e) Flow cytometry quantification of frequency shifts within intra-tumoral pro-inflammatory Ly6C^+^ (CD206^-^F4/80^+^) or MDSC-like Ly6G^+^ myeloid cells (gated on CD45^+^CD11b^+^) following indicated treatments. Quantification of MFI of M2-like macrophage marker CD206 (f) and dendritic cell marker CD11c (g) on tumoral myeloid cells following indicated treatments. For all data, N≥3 mice per group.

IHC analysis of tumors in each treatment group reflected the observed transcriptional changes which suggested a favorable pro-inflammatory TME shift allowing a reversion of T cell exclusion (**Figure 4 c**). Specifically, we found increased frequencies of intratumoral cytotoxic CD8+ T cells and a corresponding increase of granzyme-B+ (GzmB) cells following Cy pre-conditioning. We also observed increases in intratumoral PD-L1 in areas coincident with higher T cell accumulation. Interestingly, Cy did not appear to modulate Foxp3+ Treg frequencies within the tumor. Increased immune-related gene signatures in pre-conditioned tumors highlight the impact of Cy, which was further enhanced by the presence of PSCA-mCAR T cells (**Figure 4 d**). The majority of upregulated genes included markers of immune cell subsets (e.g., *Cd3e, Cd4, Cd8a, Itgam)*, monocyte/myeloid cell activation/differentiation (e.g., *Ly6c1/2, Ly6g, Csf1r, Itgam/Cd11b, Itgax/Cd11c*), T cell activation (e.g., *Gzmb, Cd274*), pro-inflammatory factors (e.g., *Il6, Nos2, IL1b, Tnf*), and T cell chemoattraction (e.g., *Ccl5, Cxcl9*). Flow cytometry analysis of pro-inflammatory monocytic Ly6C^+^ cells (gated within CD11b^+^) showed a significant increase following Cy pre-conditioning (**Figure 4 e**). Cy pre-conditioning also resulted in fewer tumor-promoting M2-like CD206^+^ myeloid cells and increased CD11c^+^ antigen presenting myeloid populations (**Figure 4 f and g**), confirming findings from the RNA sequencing analysis. Furthermore, a pro-inflammatory shift in peripheral myeloid subsets (increased peripheral Ly6C^+^ and decrease in Ly6G^+^) was observed in the blood of mice treated with the combination of Cy pre-conditioning and PSCA-mCAR T cells relative to CAR T cells alone (**Figure S7 e**). IPA network canonical pathway analysis also showed increases in innate/adaptive immune cell communication following Cy pre-conditioning, with antigen-presentation pathways most upregulated following PSCA-mCAR T cells and Cy (**Figure S7 a - d**). Overall, these data support a key role for Cy in overcoming the local immunosuppressive TME and promoting endogenous anti-tumor immunity.

### PSCA-mCAR T cells effectively target PSCA+ prostate cancer bone metastases and generates protective tumor immunity

We next assessed our therapy in a more clinically-relevant bone metastatic model. hPSCA-KI mice were i.ti. engrafted with RM9-hPSCA tumors, pre-conditioned with Cy, followed by i.v. Mock or PSCA-mCAR T cell treatment as described in **Figure 5 a**. Tumor flux imaging indicated that mice treated with Mock T cells, Mock T cells with Cy, and PSCA-mCAR T cells alone showed minimal anti-tumor responses. In contrast, PSCA-mCAR T cells in combination with Cy pre-conditioning promoted anti-tumor responses in the majority of treated mice (**Figure 5 b and S5 a**) with 50% curative responses (**Figure 5 c**). At day 7 post-CAR T cell therapy, we observed clearance of bone tumors and unremarkable gross architecture of bone marrow in the combination group (**Figure S5 b**).

**Figure 5:**
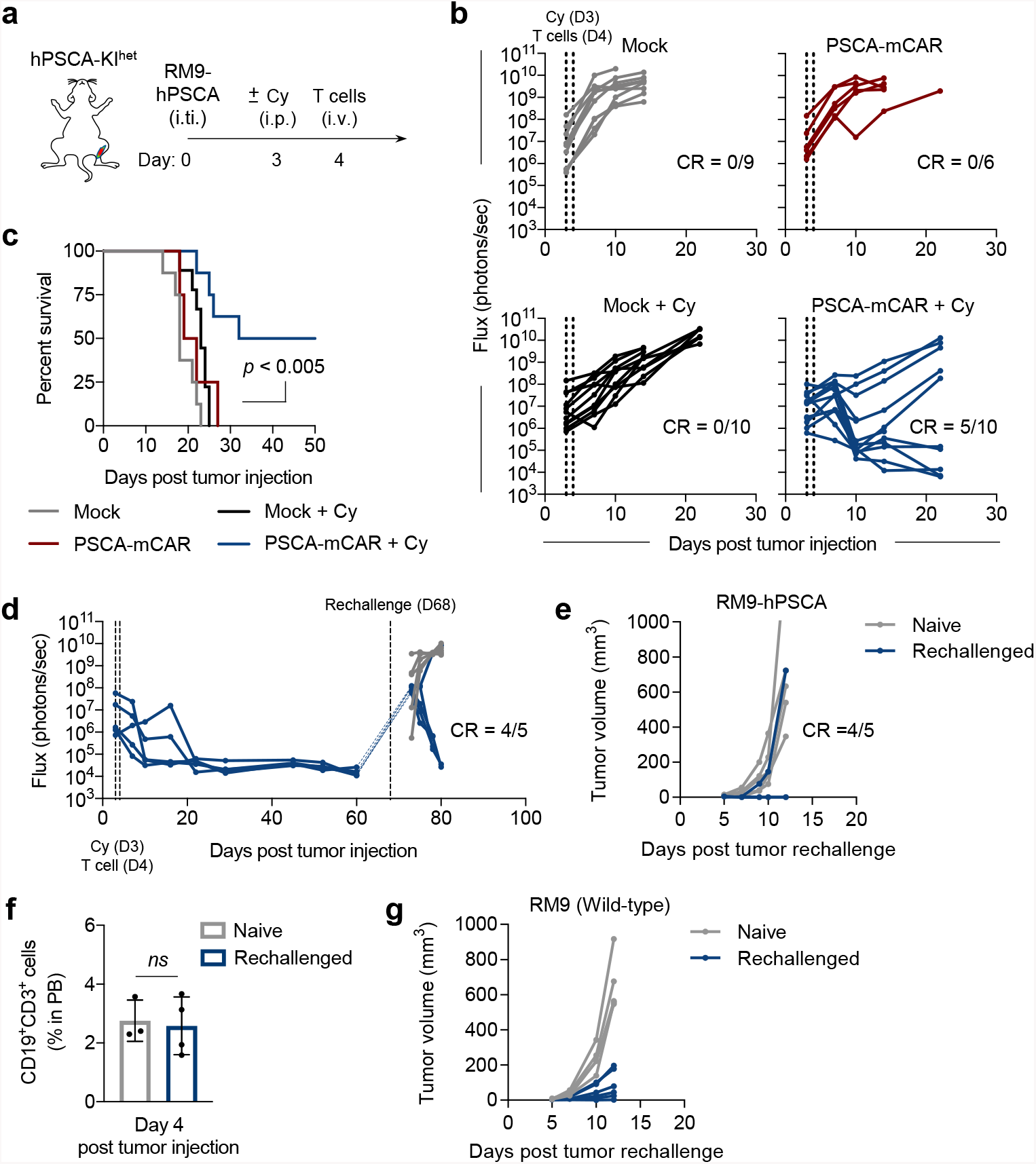
Cy pre-conditioning combined with PSCA-mCAR T cell treatment is effective *in vivo* against bone-metastatic RM9-hPSCA prostate tumors and promotes protective anti-tumor immune memory. a) Illustration of RM9-hPSCA i.ti. engrafted prostate tumors with or without Cy pre-conditioning (vehicle or Cy 100mg/kg, i.p.) 24 h prior to treatment with 5.0×10^6^ T cells (Mock or PSCA-mCAR, i.v.). b) Tumor flux in each treatment group as measured by bioluminescent imaging in each replicate mouse at indicated days post tumor injection. c) Kaplan-Meier survival for treated mice. d) Tumor flux in mice engrafted with RM9-hPSCA i.ti. which achieved CR following PSCA-mCAR treatment with Cy pre-conditioning. At day 68 post initial tumor injection, tumor naïve and previously cured mice were then rechallenged with s.c. RM9-hPSCA tumors. e) Corresponding replicate RM9-hPSCA tumor volume measurements in tumor naïve and rechallenged mice from Figure 5 d. f) Flow cytometry highlighting absence of %CD19+CD3+ (%PSCA-mCAR, gated on CD45+CD3+) in peripheral blood (PB) at 4 days post RM9-hPSCA tumor challenge in naïve or rechallenge in previously cured PSCA-mCAR + Cy mice. g) Replicate RM9 wild-type tumor volume measurements in tumor naïve and rechallenged mice previously cured by PSCA-mCAR T cell + Cy treatment. N≥5 mice per group. All data are representative of two independent experiments.

We next examined the impact on long-term anti-tumor immunity resulting from reversion of T cell exclusion observed in mice pre-conditioned with Cy and treated with CAR T cells (**Figure 4 and Figure S7**). Cured mice following Cy and CAR T cell combo treatment, along with tumor naïve control mice, were rechallenged with s.c. RM9-hPSCA tumors (**Figure 5 d, left**). Impressively, in mice rechallenged with PSCA-expressing tumor, we observed rejection in 80% of mice as shown by flux imaging and caliper tumor volume measurement (**Figure 5 d right, and 5 e**). Mice rechallenged with RM9-hPSCA failed to show a re-emergence of PSCA-mCAR T cells systemically, suggesting that tumor rejection was independent of PSCA-CAR targeting of PSCA+ tumor, and related to modified endogenous anti-tumor immunity (**Figure 5 f**). To confirm this, we repeated rechallenge in Cy and PSCA-mCAR cured mice as before, but this time rechallenged with RM9 wild-type tumors (non-PSCA, non-ffluc expressing) and again saw strong endogenous anti-tumor immunity in mice previously cured with Cy and PSCA-mCAR T cell treatment relative to tumor naïve mice (**Figure 5 g**). These data suggest that Cy and CAR T cell-mediated increases in antigen presentation, innate/adaptive immune response, and IFNγ signaling pathways results in improved primary PSCA-mCAR T cell responses, as well as a stimulation of endogenous protective immunity tumor rechallenge.

### PSCA-mCAR T cells effectively target PSCA+ KPC metastatic pancreatic cancer

In addition to its overexpression in prostate cancers and their metastases, PSCA is also highly expressed in pancreatic cancers.^17^ Accordingly, we sought to validate our Cy pre-conditioning in combination with PSCA-CAR T cells against KPC pancreatic ductal adenocarcinoma (PDAC) tumor cells engineered to express hPSCA (KPC-hPSCA). Importantly, KPC tumors harbor features of PDAC, including intraepithelial neoplasia and a robust inflammatory reaction including exclusion of effector T cells.^18^ KPC-hPSCA were administered i.v., which induces lung metastasis, and were pre-conditioned with Cy, followed with i.v. Mock or PSCA-mCAR T cell treatment as indicated in **Figure 6 a**. Quantification of tumor flux post T cell treatment showed that mice treated with Mock T cells alone or Mock T cells with Cy exhibited minimal anti-tumor responses (**Figure 6 b and Figure S6 a**) as expected. Interestingly, PSCA-mCAR T cells treatment alone in this model showed pronounced but transient anti-tumor activity, distinct from the prostate tumor model. Impressively, the combination of Cy pre-conditioning and PSCA-mCAR T cells showed a more potent anti-tumor activity resulting in extended survival and curative responses in over 60% of mice (**Figure 6 c**). IHC of lung tissues at 13 days post-T cell treatment showed disease-free mice treated with PSCA-mCAR T cells and Cy (**Figure S6 b**). Combined, these models demonstrate the utility of pre-conditioning with Cy to improve PSCA-CAR T cell therapy responses in solid tumors.

**Figure 6:**
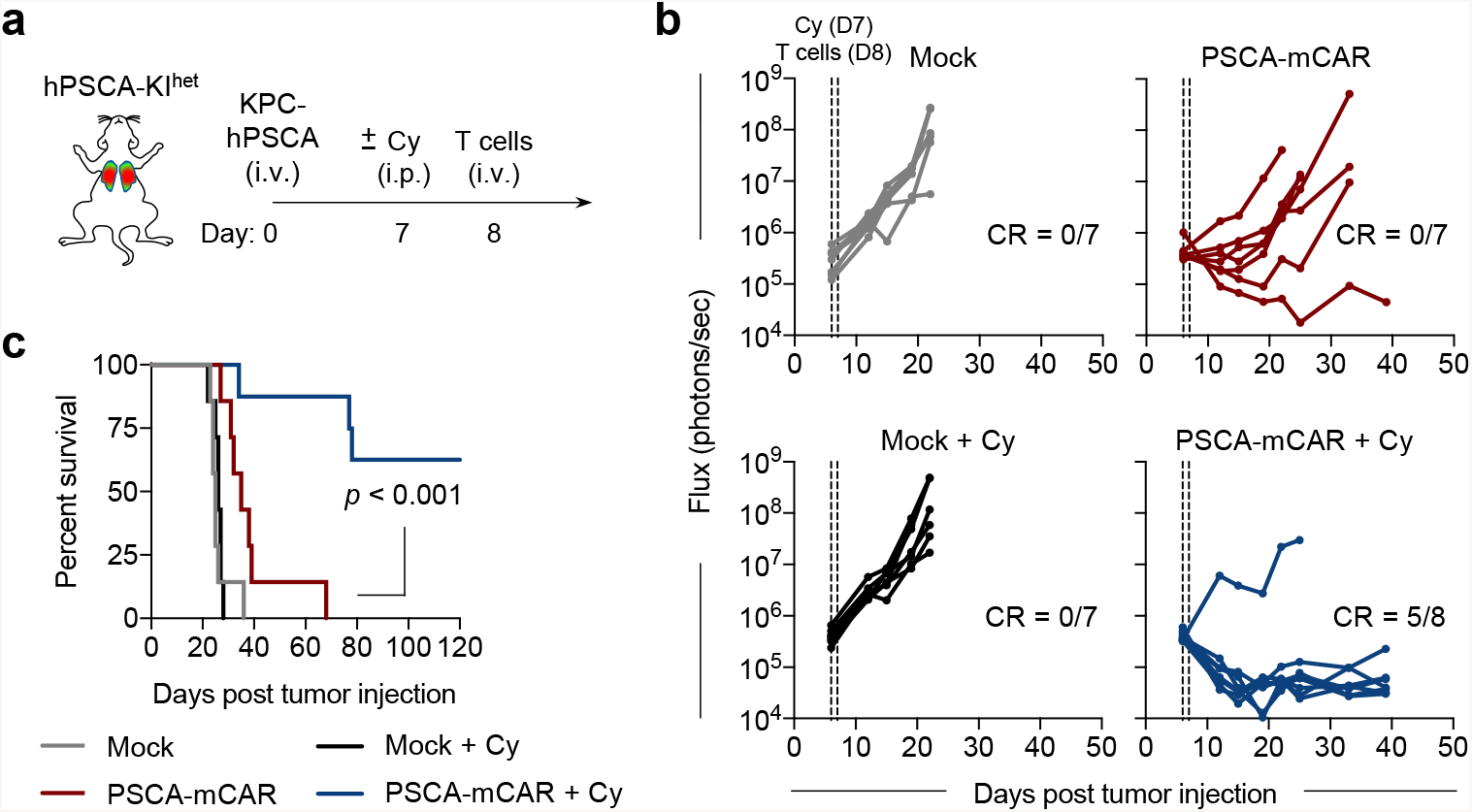
Cy pre-conditioning combined with PSCA-mCAR T cell treatment is effective *in vivo* against PSCA+ pancreatic tumor models. a) Illustration of KPC-hPSCA i.v. engrafted pancreatic tumors established in the lungs, with or without Cy pre-conditioning (vehicle or Cy 100mg/kg, i.p.) 24 h prior to treatment with 5.0×10^6^ T cells (Mock or PSCA-mCAR, i.v.). b) Tumor flux in each treatment group as measured by bioluminescent imaging in each replicate mouse at indicated days post tumor injection. c) Kaplan-Meier survival plot for mice in each group indicated. N≥7 mice per group. All data are representative of two independent experiments.

## Discussion

In this study, we investigated the safety and efficacy of PSCA CAR T cell therapy using immunocompetent hPSCA-KI mice. We discovered that robust local intratumoral activity, proliferation and therapeutic responses of PSCA-CAR T cells were dependent on pre-conditioning with Cy. PSCA-CAR T cell therapy demonstrated safety in mice with physiological hPSCA expression in normal tissues, mimicking patterns of expression found in humans.^15,16^ We showed a lack of therapeutic responses with PSCA-CAR T cell treatment alone in this system, contrasting with the complete responses previously observed using human xenograft tumor models in immunocompromised NSG mice.^10^ The dramatic improvement of CAR T cell activity and survival of mice following Cy pre-conditioning led us to postulate that in addition to its conventional lymphodepleting effects, Cy also mediates substantial modulation of the solid tumor TME. Accordingly, we showed that Cy pre-conditioning resulted in rapid CAR and endogenous T cell infiltration within the tumor, re-shaped the TME resulting in shifts in myeloid cells from M2 to M1 phenotypes, increased endogenous antigen presentation pathways, and increased innate/adaptive immune response signaling, all further enhanced by combination with CAR T cells. Importantly, we also showed CAR antigen-independent protective immunity against tumor rechallenge in this model. Together, these data emphasize three important features: 1) commonly used immunocompromised xenograft animal models greatly underrepresent the impact of the host immune system thereby overestimating CAR T cell activity, 2) independent of peripheral lymphodepleting effects, Cy modulates the local TME and may be critical to unleashing the full potential of solid tumor CAR T cell therapies, and 3) the combination of Cy pre-conditioning along with CAR therapy in our model also modifies host immunity to elicit protection against tumor rechallenge.

Prior studies have demonstrated Cy-mediated depletion of peripheral T cells, particularly Tregs, may act as a cytokine sink to alter CD8 T cell activity.^7,19,20^ Studies suggest that Cy-mediated depletion of Tregs is highly transient, and modulation of dendritic cell function within the TME may explain improved anti-tumor immunity.^21^ Similarly, we did not observe striking changes in tumoral Foxp3+ Tregs following Cy in our model, suggesting alternative modifications of the local TME. We found that as early as 24 h following PSCA-mCAR T cell treatment, clear intratumoral trafficking and accumulation of CAR T cells were observed in Cy pre-conditioned mice. This robust tumoral recruitment of PSCA-mCAR T cells was observed prior to the peak peripheral lymphodepleting effects of Cy in this model. Once maximal lymphodepletion was achieved, 3 days following T cell treatment, we then observed great increases in intratumoral T cell expansion both in antigen specific PSCA-mCAR, and to a lesser degree in adoptively transferred non-targeting Mock T cells. This contrasts with previous findings that indicate intratumoral T cell homing or migration aided by Cy is reserved for antigen-specific T cells.^22^ In addition to the direct T cell modulating effects of Cy, pre-conditioning lymphodepletion enhances IL-2, IL-7, and IL-15 cytokine signaling to improve systemic persistence of CD19-CAR T cells.^22,23^ However, we found no increases in peripheral CAR T cells despite robust local anti-tumor responses. Additionally, prior to maximal peripheral lymphodepletion, Cy modified the local TME, improving infiltration of endogenous T cells, IFNγ signaling in the TME, as well as an M2-like to M1-like macrophage shift. This result is consistent with prior work showing pro-inflammatory modulation by Cy in the TME, which may synergize with type-I IFN cytokines, which we observe especially in combination with antigen specific PSCA-mCAR T cells.^24,25^

Cy pre-conditioning greatly enhanced PSCA-mCAR T cell activity in the immunocompetent models we tested, but did not demonstrate curative responses in all treated mice. This implies that additional CAR T cell resistance mechanisms may limit maximal therapeutic benefits of this approach and must be investigated further. Major resistance mechanisms include hampered T cell trafficking or persistence in tumors, intrinsic and/or acquired resistance to CAR T cell therapy driven in part by immune suppressive pathways, and tumor antigen escape from single-antigen targeted CAR T cell strategies. With regards to trafficking, it is possible that a lack of immediate antigen engagement by systemically-administered CAR may limit peripheral survival and trafficking to solid tumors as there is observed trapping of adoptively-transferred T cells in first-pass tissues, including lung and liver.^26^ Interestingly, PSCA-mCAR T cells, both with and without pre-conditioning, demonstrated greater therapeutic activity in the lung metastatic KPC-PDAC model, perhaps taking advantage of earlier lung trafficking of systemically administered T cells. A recent study in a melanoma model showed that lymphodepleting chemotherapy pre-conditioning promoted peripheral immunosuppressive MDSCs, which limited persistence of tumor infiltrating T cells and resulted in greater disease progression.^27^ While at later timepoints, Cy-mediated immunosuppression may contribute to tumor recurrences in our models, our data strongly support the early benefits of Cy pre-conditioning in re-shaping the tumor microenvironment to promote CAR T cell responses.

Further studies are warranted to determine the mechanism(s) responsible for the protective immunity following Cy and PSCA-CAR T cell therapy. Immune checkpoint pathways may also limit therapeutic responses and have been highlighted in multiple preclinical and clinical CAR T cell programs.^28,29^ In support of this acquired tumor resistance mechanism, we observed robust *in vitro* and *in vivo* induction of PD-1/PDL1 signaling following PSCA-CAR T cell treatment.^30-32^ Future studies will investigate the benefits of combining PSCA-CAR T cells with immune checkpoint blockade and other acquired or intrinsic immunosuppressive pathways. Antigen escape mechanisms are a major challenge in achieving durable responses to CAR T cell therapy in both hematological and solid tumor malignancies.^33-35^ In our model, tumor cells were clonally positive or FACS sorted for uniformly positive PSCA expression, which would greatly reduce the possibility of an antigen-negative outgrowth following CAR T cell therapy. However, transcriptional downregulation or novel PSCA mutations in response to CAR T cell pressure may have occurred, similar to clinical observations following CD19-CAR T cell therapy.^33,36^ Prior clinical experience with hematological malignancies and solid tumors alike suggests antigen heterogeneity is likely a key barrier that will require multi-targeted CAR T cell approaches.

To our knowledge, this is the first example of a preclinical model to assess the safety of PSCA-targeted agents, including CAR T cell therapy. Several groups have developed immunocompetent mouse models to evaluate the safety of T cell therapeutic approaches, including those targeting CD19, CEA, and HER2.^14,37,38^ In pre-clinical CD19-CAR T cell models, the severity of toxicity (i.e. CRS, neurotoxicity, and in some cases death) was increased in the presence of lymphodepleting pre-conditioning regimens.^39^ Further, in human CEA transgenic mouse models, it was found that CEA-targeted CAR T cells were only effective when combined with lymphodepletion. In these mice, toxicities (colitis, weight loss, liver toxicity, and death) were only seen in mice pre-conditioned with total body irradiation (TBI) alone or a triple combination of fludarabine, Cy, and TBI.^14,40^ In contrast, tumor bearing hPSCA-KI mice who received Cy-preconditioning and PSCA-CAR T cell therapy displayed no overt toxicities, including normal tissues expressing PSCA; however, expression intensity and pattern of PSCA in our hPSCA-KI model may not completely represent human biology. Our phase 1 clinical trial (NCT03873805) is evaluating the safety, feasibility, and biological activity of PSCA-CAR T cells in patients with metastatic castration-resistant prostate cancer, and based on these preclinical studies we have added a lymphodepletion arm to assess the effects on the TME and clinical outcomes.

## Materials and Methods

### Cell Lines

The Ras/Myc transformed prostate cancer line, RM9 and firefly-luciferase expressing LSL-Kras^G12D/+^; LSL-Trp53^R172H/+^; Pdx-1-Cre (KPC) cell lines were a kind gift from Dr. Timothy C. Thompson at MD Anderson Cancer Center Dr. Edwin Manuel at City of Hope respectively.^41,42^ Cell lines were cultured in Dulbecco’s Modified Eagles Medium (Life Technologies), containing 10% fetal bovine serum (FBS, Hyclone), 25 mM HEPES (Irvine Scientific), 2 mM L-Glutamine (Fisher Scientific), and 1X antibiotic-antimycotic (1X AA, Gibco) (cDMEM). PLAT-E retroviral packaging cells (Cell Biolabs Inc.) were cultured in cDMEM with addition of 1 μg/mL puromycin and 10 μg/mL blasticidin (InvivoGen) prior to transfection. All cells were cultured at 37°C with 5% CO2.

### Animals

Animal experiments were performed under protocols approved by the City of Hope Institutional Animal Care and Use Committee. For all *in vivo* studies, 6–8 week-old male mice were used. Human prostate stem-cell antigen (PSCA) knock-in (hPSCA-KI) C57BL/6j mice, whose generation has been described previously, were provided by Dr. Robert E. Reiter and Dr. Anna Wu at UCLA.^15^ To prevent possible rejection of tumor lines that endogenously express murine PSCA, homozygous hPSCA-KI mice were F1 crossed with wild-type C57BL/6j mice to generate heterozygous hPSCA-KI mice. Unless otherwise indicated, all mice used in experiments were heterozygous hPSCA-KI. For experiments in immunocompromised animals, 6 to 8 week-old RAG2^-/-^ (Jackson Laboratories) mice were used.

### Genotyping PCR

To confirm the genotype of hPSCA-KI mice, genomic DNA was isolated and analyzed by PCR. Genomic DNA was collected from tail or ear tissue isolated via Qiagen DNA Mini Kit (Qiagen) according to manufacturer’s protocol. Isolated DNA samples were analyzed in independent master mixes containing primers, Accustart PCR SuperMix (Quantabio), and sterile DNAse/RNAse-free water. Three primer sequences (kindly provided by Dr. Robert Reiter at UCLA) were used to detect murine genomic PSCA DNA (from 5’ and 3’ arms) and hPSCA cDNA. mPSCA-5’arm primer sequence: TGTCACTGTTGACTGTGGGTAGCA, mPSCA-3’arm primer sequence: CTTACTTGATAGGAGGGCTCAGCA, hPSCA primer sequence: CCAGAGCAGCAGGCCGAGTGCA. PCR was performed on a FlexCycler^2^ (Analytik Jena) programmed for 94°C for 1 minute, 35 cycles at (94°C for 20 seconds, 59°C for 20 seconds, 72°C for 45 seconds), and a final run at 72°C for 3 minutes. PCR products were loaded on to 2% agarose gels, run at 150 V for approximately 20 minutes, and stained in SybrSafe (Life Technologies) prior to imaging on a BioSpectrum MultiSpectral Imaging System (UVP).

### DNA Constructs, Tumor Lentiviral Transduction and Retrovirus Production

Tumor cells were engineered to express firefly-luciferase (ffluc) and tumor antigen human PSCA (hPSCA) by sequential transduction with epHIV7 lentivirus carrying the ffluc gene under the control of the EF1α promoter, and epHIV7 lentivirus carrying the human PSCA gene under the control of the EF1α promoter as described previously.^10^ The scFv sequence from the murine anti-human PSCA antibody clone (1G8) was used to develop the murine CAR (mCAR) construct.^43^ The extracellular spacer domain included the murine CD8 hinge region followed by a murine CD8 transmembrane domain.^44,45^ The intracellular co-stimulatory signaling domain contained the murine 4-1BB followed by a murine CD3ζ cytolytic domain as previously described.^46^ The CAR sequence was separated from a truncated murine CD19 gene (mCD19t) by a T2A ribosomal skip sequence and cloned into the pMYs retrovirus backbone (Cell Biolabs Inc). Production of retrovirus used to transduce primary murine T cells were performed as previously described.^47^ The CAR sequence was separated from a truncated murine CD19 gene (mCD19t) by a T2A ribosomal skip sequence and cloned into the pMYs retrovirus backbone under the control of a hybrid MMLV/MSCV promoter (Cell Biolabs Inc). The ffluc sequence was also cloned into the pMYs retrovirus backbone. Retrovirus was produced by transfecting the ecotropic retroviral packaging cell line, PLAT-E, with addition of PSCA-mCAR or ffluc retrovirus backbone plasmid DNA using FuGENE HD transfection reagent (Promega). Viral supernatants were collected after 24, 36, and 48 h, pooled, and stored at −80ºC in aliquots for future T cell transductions.

### Murine T Cell Isolation, Transduction, and *Ex Vivo* Expansion

Splenocytes were obtained by manual digestion of spleens from male heterozygous hPSCA-KI mice. Enrichment of T cells was performed by EasySep™ mouse T cell isolation kit per manufacturer’s protocol (StemCell Technologies). Retroviral transduction with PSCA-mCAR and/or ffluc and subsequent expansion were performed as previously described.^47^

### Flow Cytometry

Flow cytometric analysis was performed as previously described.^10,47^ Briefly, cells were resuspended in Hank’s balanced salt solution without Ca^2+^, Mg^2+^, or phenol red (HBSS^-/-^, Life Technologies) containing 2% FBS and 1x Antibiotic-Antimycotic (AA, Gibco) (FACS buffer). Single cell suspensions from mouse tissues or tumors were incubated for 15 minutes at ambient room temperature with rat anti-mouse Fc Block™ (BD Pharmingen). Cells were then incubated with primary antibodies for 30 minutes at 4ºC in the dark. For secondary staining, cells were washed twice prior to 30 minutes incubation at 4ºC in the dark with either Brilliant Violet 510 (BV510), fluorescein isothiocyanate (FITC), phycoerythrin (PE), peridinin chlorophyll protein complex (PerCP), PerCP-Cy5.5, PE-Cy7, allophycocyanin (APC), or APC-Cy7 (or APC-eFluor780)-conjugated antibodies. Anti-mouse antibodies against mCD206 (Biolegend, Clone: C068C2), mLy6G (Biolegend, Clone: 1A8), mLy6C (Biolegend, Clone: HK1.4), mCD11b (Biolegend, Clone: M1/70), mCD11c (Biolegend, Clone: N418), mMHC Class II (I-A/I-E, Biolegend, Clone: M5/144.15.2), mCD274 (mPD-L1, Biolegend, Clone:10F.9G2), mCD45 (Biolegend, Clone: 30-F11), mCD19 (BD Biosciences, Clone: 1D3), mCD3 (Biolegend, Clone: 17A2), mCD279 (mPD-1, Biolegend, Clone: 29F.1A12), mCD4 (Biolegend, Clone: RM4-5), mCD137 (ThermoFisher, Clone: 17b5), mCD8a (Miltenyi Biotec, Clone: 53-6.7), and goat anti-mouse Ig (BD Biosciences) were used for flow cytometry. Additionally, a mouse anti-human PSCA antibody (clone 1G8) used for flow cytometry was from the laboratory of Dr. Robert Reiter at UCLA ^16^. Cell viability was determined using 4’, 6-diamidino-2-phenylindole (DAPI, Sigma). Flow cytometry was performed on a MACSQuant Analyzer 10 (Miltenyi Biotec), and data was analyzed with FlowJo software (v10, TreeStar).

### *In Vitro* Tumor Killing and T Cell Functional Assays

For tumor killing assays, PSCA-mCAR T cells and tumor targets were co-cultured at indicated effector:tumor (E:T) ratios in the absence of exogenous cytokines or antibiotics in 96-well plates for 24 to 72 h and analyzed by flow cytometry. Tumor killing was calculated by comparing DAPI-negative (viable) mCD45-negative cell counts in co-cultures with CAR T cells relative to that observed in co-culture with Mock (untransduced) T cells. T cell functional analysis from these assays were also determined by flow cytometry, using indicated cell surface antibodies.

### ELISA Cytokine Assays

Cell-free supernatants from tumor killing assays were collected at indicated times and frozen at − 20°C for further analysis. Murine IFNγ and IL-2 cytokines were measured in supernatants according to the Murine IFNγ and IL-2 ELISA Ready-SET-GO!® ELISA kit (Invitrogen) according to manufacturer’s protocol, respectively. Plates were read at 450 nm using a Cytation3 imaging reader with Gen5 microplate software v3.05 (BioTek).

### Tumor RNA Isolation, Sequencing, and Analysis

Tumor tissue samples harvested from mice were immediately snap-frozen in liquid nitrogen and stored at −80°C for further processing. Frozen tumor samples were placed in Green RINO RNA lysis tubes for bead homogenization in a Bullet Blender homogenizer (Next Advance) in the presence of TRIzol at 4°C. Lysate was then chloroform extracted for RNA and purified via a Qiagen RNA Easy isolation kit (Qiagen) according to manufacturer’s protocol. Libraries for stranded polyA RNA-Seq were created using the KAPA mRNA HyperPrep kit (Roche).

Sequencing of 51 bp single-end reads was performed using a HiSeq2500 regular run. Base calling (de-multiplexing samples between and within labs by 6 bp barcodes, from a 7 bp index read) was performed using bcl2fastq v2.18. Reads were aligned against the mouse genome (mm10) using TopHat2 ^48^. Read counts were tabulated using htseq-count,^49^ with UCSC known gene annotations (TxDb.Mmusculus.UCSC.mm10.knownGene, downloaded 8/30/2018).^50^ Fold-change values were calculated from Fragments Per Kilobase per Million reads (FPKM) normalized expression values, which were also used for visualization (following a log2 transformation).^51^ Aligned reads were counted using GenomicRanges.^52^ Scripts are a modified version of a template for RNA-Seq gene expression analysis (https://github.com/cwarden45/RNAseq_templates/tree/master/TopHat_Workflow) and RNA-Seq data for this study was deposited in X. Expression of log2(FPKM + 0.1) expression was visualized in heatmaps using heatmap.3 (https://github.com/obigriffith/biostar-tutorials/blob/master/Heatmaps/heatmap.3.R). Custom gene sets defined in Ingenuity Pathway Analysis (IPA, Ingenuity® Systems, www.ingenuity.com) was used for generation of specific heatmap gene sets used in the current study. Candidate genes were selected by pair-wise comparisons between treatment groups, looking for genes with a |fold-change| > 1.5. Genes were filtered to only include transcripts with an FPKM expression level of 1 (after a rounded log2-transformation) in at least 50% of samples as well as genes that are greater than 150 bp. An expression level of FPKM greater than 1 was used to reduce false positives. Expression level FPKM data was then used to generated 1 vs 1 treatment group comparisons. IPA pathway analysis was used to calculate canonical pathway enrichments among treatment groups and generation of heatmaps using the provided set of genes. Gene Set Enrichment Analysis (GSEA) was run on log2(FPKM + 0.1) expression values, with up-regulated enrichment results for Gene Ontology (GO) Biological Process categories in MSigDB.^53-55^

### *In Vivo* Tumor Studies

For *in vivo* intratibial (i.ti.) tumor studies, 2.5 × 10^4^ RM9-hPSCA/ffluc cells were prepared HBSS^-/-^ and i.ti. engrafted in 6–8 week-old male heterozygous hPSCA-KI mice as described previously.^10^ For *in vivo* subcutaneous (s.c.) tumor studies, 0.2 − 1.0 ×10^6^ RM9-hPSCA/ffluc cells were prepared in HBSS^-/-^ and s.c. engrafted in 6–8 week-old male heterozygous hPSCA-KI mice. Tumor growth was monitored at least twice per week via biophotonic imaging (LagoX, Spectral Instruments) or by volumetric caliper measurement (length x width x height = mm^3^). Flux signals were analyzed by noninvasive optimal imaging as previously described.^10,47^ For i.ti. studies, on day 3 or 4 post engraftment, mice were grouped based on flux signal and i.p. injected with the indicated dose of cyclophosphamide (Cy, Sigma-Aldrich) or vehicle (sterile PBS). For s.c. studies, mice were treated i.p. with Cy or vehicle once tumor volumes reached an average of 100 mm^3^. For all studies, mice received intravenous (i.v.) treatment with 5.0 × 10^6^ Mock or PSCA-mCAR T cells. Flux imaging or caliper measurement continued until scheduled harvest or humane endpoints were reached. For tumor rechallenge studies, tumor burden was measured by flux imaging until complete responses (CR) were achieved and maintained for at least 30 days. Mice achieving a CR, and age matched tumor naïve hPSCA-KI control mice, were then rechallenged by s.c. injection of 0.5 to 1.0 x 10^5^ RM9-hPSCA/ffluc or RM9 wild-type (non-PSCA, non-ffluc) cells and engraftment was measured by flux imaging and/or volumetric caliper measurement. Humane endpoints were used in determining survival. Mice were euthanized upon reaching i.ti. and s.c. tumor volumes in excess of 1500 mm^3^, showing signs of distress, labored or difficulty breathing, weight loss, impaired mobility, or evidence of being moribund.

Peripheral blood (PB) was collected from isoflurane-anesthetized mice by retro-orbital (RO) bleed through heparinized capillary tubes (Chase Scientific) into polystyrene tubes containing a heparin/PBS solution (1000 U/mL, Sagent Pharmaceuticals) at the indicated time points. Volume of each RO blood draw was recorded for cell quantification per µL blood. Tumor and normal tissue samples harvested from euthanized mice were collected into ice-cold PBS and subsequently processed using a murine tumor digestion kit (Miltenyi Biotec) according to manufacturer’s protocol. Red blood cells (RBCs) from either RO bleed or tissue collection were lysed with 1X Pharmlyse buffer (BD Biosciences) according to manufacturer’s protocol and then washed, stained, and analyzed by flow cytometry.

## Immunohistochemistry and RNA *in situ* Hybridization

Tumor tissue was fixed in 4% paraformaldehyde (4% PFA, Boston BioProducts) and stored in 70% ethanol until further processing. Immunohistochemistry was performed by the Research Pathology Core at City of Hope. Briefly, paraffin sections (5 µm) were stained with hematoxylin & eosin (H&E, Sigma-Aldrich), mouse anti-human PSCA antibody (Abnova, H00008000-M03, Clone: 5C2), anti-mouse CD3 (Abcam, ab16669, Clone: SP7), anti-mouse granzyme-B (Abcam, ab4059, polyclonal), anti-mouse CD8a (Cell Signaling, 98941S, Clone: D4W2Z), anti-mouse PD-L1 (Abcam, AB80276, Clone: MIH6), anti-mouse FOXP3 (Abcam, Clone: EPR22102-37) according to manufacturer’s protocol. RNA *in situ* hybridization of human PSCA was performed by RNAScope™ (ACD). Images were obtained using the Nanozoomer 2.0HT digital slide scanner and the associated NDP.view2 software (Hamamatsu).

## Statistical Analysis

Data are presented as mean ± standard error of the mean (SEM), unless otherwise stated. Statistical comparisons between groups were performed using the unpaired two-tailed Student’s *t* test to calculate *p* value, unless otherwise stated. \**p* < 0.05, **p < 0.01, ***p < 0.001; NS, not significant.

## Supplemental Information

Supplemental figures and legends are available and included as separate document.

## Acknowledgments

We thank staff members of the Animal Facility and Small Animal Imaging Core, Pathology Core, Integrative Genomics and Bioinformatics Core in the Beckman Research Institute at the City of Hope Comprehensive Cancer Center for excellent technical assistance, supported by the National Cancer Institute of the National Institutes of Health under award number P30CA033572. We thank Charles Warden and Dr. Xiwei Wu of the Integrative Genomics Core for their technical assistance in RNA sequencing analysis. We would also like to thank Dr. Sandra Thomas at City of Hope for contributions to editing of manuscript. The content is solely the responsibility of the authors and does not necessarily represent the official views of the National Institutes of Health.

## Availability of Data

All data is available upon reasonable request.

## Author Contributions

JPM and SJP conceived and designed the study and wrote the manuscript. JPM performed majority of experiments and data analyses. DT, AKP, KTK, LSL, HJL, BJG, and WC contributed to in vivo experiment execution, cloning, and/or data collection. RER and CPT provided access to transgenic mice and were consulted on related studies. CM, AMW, RER, TBD, and SJF were consulted on methods used and/or assisted in writing the manuscript.

## References

1. Wang, Z, Chen, W, Zhang, X, Cai, Z, and Huang, W (2019). A long way to the battlefront: CAR T cell therapy against solid cancers. J Cancer 10: 3112–3123.

2. Priceman, SJ, Forman, SJ, and Brown, CE (2015). Smart CARs engineered for cancer immunotherapy. Curr Opin Oncol 27: 466–474.

3. Mardiana, S, Solomon, BJ, Darcy, PK, and Beavis, PA (2019). Supercharging adoptive T cell therapy to overcome solid tumor-induced immunosuppression. Sci Transl Med 11.

4. Bonaventura, P, Shekarian, T, Alcazer, V, Valladeau-Guilemond, J, Valsesia-Wittmann, S, Amigorena, S, et al. (2019). Cold Tumors: A Therapeutic Challenge for Immunotherapy. Front Immunol 10: 168.

5. Fleming, V, Hu, X, Weber, R, Nagibin, V, Groth, C, Altevogt, P, et al. (2018). Targeting Myeloid-Derived Suppressor Cells to Bypass Tumor-Induced Immunosuppression. Front Immunol 9: 398.

6. Muranski, P, Boni, A, Wrzesinski, C, Citrin, DE, Rosenberg, SA, Childs, R, et al. (2006). Increased intensity lymphodepletion and adoptive immunotherapy--how far can we go? Nature clinical practice Oncology 3: 668–681.

7. Gattinoni, L, Finkelstein, SE, Klebanoff, CA, Antony, PA, Palmer, DC, Spiess, PJ, et al. (2005). Removal of homeostatic cytokine sinks by lymphodepletion enhances the efficacy of adoptively transferred tumor-specific CD8+ T cells. J Exp Med 202: 907–912.

8. Mikyskova, R, Indrova, M, Pollakova, V, Bieblova, J, Simova, J, and Reinis, M (2012). Cyclophosphamide-induced myeloid-derived suppressor cell population is immunosuppressive but not identical to myeloid-derived suppressor cells induced by growing TC-1 tumors. Journal of immunotherapy (Hagerstown, Md: 1997) 35: 374–384.

9. Bracci, L, Schiavoni, G, Sistigu, A, and Belardelli, F (2014). Immune-based mechanisms of cytotoxic chemotherapy: implications for the design of novel and rationale-based combined treatments against cancer. Cell death and differentiation 21: 15–25.

10. Priceman, SJ, Gerdts, EA, Tilakawardane, D, Kennewick, KT, Murad, JP, Park, AK, et al. (2018). Co-stimulatory signaling determines tumor antigen sensitivity and persistence of CAR T cells targeting PSCA+ metastatic prostate cancer. Oncoimmunology 7: e1380764.

11. Gu, Z, Thomas, G, Yamashiro, J, Shintaku, IP, Dorey, F, Raitano, A, et al. (2000). Prostate stem cell antigen (PSCA) expression increases with high gleason score, advanced stage and bone metastasis in prostate cancer. Oncogene 19: 1288–1296.

12. Reiter, RE, Gu, Z, Watabe, T, Thomas, G, Szigeti, K, Davis, E, et al. (1998). Prostate stem cell antigen: A cell surface marker overexpressed in prostate?cancer. Proceedings of the National Academy of Sciences 95: 1735–1740.

13. Semenkow, S, Li, S, Kahlert, UD, Raabe, EH, Xu, J, Arnold, A, et al. (2017). An immunocompetent mouse model of human glioblastoma. Oncotarget 8: 61072–61082.

14. Wang, L, Ma, N, Okamoto, S, Amaishi, Y, Sato, E, Seo, N, et al. (2016). Efficient tumor regression by adoptively transferred CEA-specific CAR-T cells associated with symptoms of mild cytokine release syndrome. Oncoimmunology 5: e1211218–e1211218.

15. Zhang, M, Kobayashi, N, Zettlitz, KA, Kono, EA, Yamashiro, JM, Tsai, W-TK, et al. (2019). Near-Infrared Dye-Labeled Anti-Prostate Stem Cell Antigen Minibody Enables Real-Time Fluorescence Imaging and Targeted Surgery in Translational Mouse Models. Clin Cancer Res 25: 188–200.

16. Gu, Z, Yamashiro, J, Kono, E, and Reiter, RE (2005). Anti-prostate stem cell antigen monoclonal antibody 1G8 induces cell death in vitro and inhibits tumor growth in vivo via a Fc-independent mechanism. Cancer Res 65: 9495–9500.

17. Katari, UL, Keirnan, JM, Worth, AC, Hodges, SE, Leen, AM, Fisher, WE, et al. (2011). Engineered T cells for pancreatic cancer treatment. HPB (Oxford) 13: 643–650.

18. Lee, JW, Komar, CA, Bengsch, F, Graham, K, and Beatty, GL (2016). Genetically Engineered Mouse Models of Pancreatic Cancer: The KPC Model (LSL-Kras(G12D/+);LSL-Trp53(R172H/+); Pdx-1-Cre), Its Variants, and Their Application in Immuno-oncology Drug Discovery. Curr Protoc Pharmacol 73: 14.39.11-14.39.20.

19. Scurr, M, Pembroke, T, Bloom, A, Roberts, D, Thomson, A, Smart, K, et al. (2017). Low-Dose Cyclophosphamide Induces Antitumor T-Cell Responses, which Associate with Survival in Metastatic Colorectal Cancer. Clin Cancer Res 23: 6771–6780.

20. Lutsiak, MEC, Semnani, RT, De Pascalis, R, Kashmiri, SVS, Schlom, J, and Sabzevari, H (2005). Inhibition of CD4+25+ T regulatory cell function implicated in enhanced immune response by low-dose cyclophosphamide. Blood 105: 2862–2868.

21. Radojcic, V, Bezak, KB, Skarica, M, Pletneva, MA, Yoshimura, K, Schulick, RD, et al. (2010). Cyclophosphamide resets dendritic cell homeostasis and enhances antitumor immunity through effects that extend beyond regulatory T cell elimination. Cancer Immunology, Immunotherapy 59: 137–148.

22. Bracci, L, Moschella, F, Sestili, P, La Sorsa, V. Valentini, M, Canini, I, et al. (2007). Cyclophosphamide Enhances the Antitumor Efficacy of Adoptively Transferred Immune Cells through the Induction of Cytokine Expression, B-Cell and T-Cell Homeostatic Proliferation, and Specific Tumor Infiltration. Clinical Cancer Research 13: 644.

23. Ninomiya, S, Narala, N, Huye, L, Yagyu, S, Savoldo, B, Dotti, G, et al. (2015). Tumor indoleamine 2,3-dioxygenase (IDO) inhibits CD19-CAR T cells and is downregulated by lymphodepleting drugs. Blood 125: 3905–3916.

24. Moschella, F, Torelli, GF, Valentini, M, Urbani, F, Buccione, C, Petrucci, MT, et al. (2013). Cyclophosphamide induces a type I interferon-associated sterile inflammatory response signature in cancer patients’ blood cells: implications for cancer chemoimmunotherapy. Clin Cancer Res 19: 4249–4261.

25. Schiavoni, G, Sistigu, A, Valentini, M, Mattei, F, Sestili, P, Spadaro, F, et al. (2011). Cyclophosphamide synergizes with type I interferons through systemic dendritic cell reactivation and induction of immunogenic tumor apoptosis. Cancer Res 71: 768–778.

26. Visioni, A, Kim, M, Wilfong, C, Blum, A, Powers, C, Fisher, D, et al. (2018). Intra-arterial Versus Intravenous Adoptive Cell Therapy in a Mouse Tumor Model. Journal of immunotherapy (Hagerstown, Md: 1997) 41: 313–318.

27. Innamarato, P, Kodumudi, K, Asby, S, Schachner, B, Hall, M, Mackay, A, et al. (2020). Reactive Myelopoiesis Triggered by Lymphodepleting Chemotherapy Limits the Efficacy of Adoptive T Cell Therapy. Molecular Therapy 28: 2252–2270.

28. Bonaventura, P, Shekarian, T, Alcazer, V, Valladeau-Guilemond, J, Valsesia-Wittmann, S, Amigorena, S, et al. (2019). Cold Tumors: A Therapeutic Challenge for Immunotherapy. Frontiers in Immunology 10.

29. Elia, AR, Caputo, S, and Bellone, M (2018). Immune Checkpoint-Mediated Interactions Between Cancer and Immune Cells in Prostate Adenocarcinoma and Melanoma. Frontiers in Immunology 9.

30. Pistillo, MP, Carosio, R, Banelli, B, Morabito, A, Mastracci, L, Ferro, P, et al. (2019). IFN-γ upregulates membranous and soluble PD-L1 in mesothelioma cells: potential implications for the clinical response to PD-1/PD-L1 blockade. Cellular & Molecular Immunology.

31. Basham, JH, and Geiger, TL (2016). Opposing Effects of PD-1/PD-L1/L2 Engagement and IFN-γ/TNF-α in the Treatment of AML w/ Anti-CD33 Chimeric Antigen Receptor-Modified T Cells. Blood 128: 5891–5891.

32. Wei, J, Luo, C, Wang, Y, Guo, Y, Dai, H, Tong, C, et al. (2019). PD-1 silencing impairs the anti-tumor function of chimeric antigen receptor modified T cells by inhibiting proliferation activity. Journal for ImmunoTherapy of Cancer 7: 209.

33. Majzner, RG, and Mackall, CL (2018). Tumor Antigen Escape from CAR T-cell Therapy. Cancer Discovery 8: 1219.

34. Hegde, M, Mukherjee, M, Grada, Z, Pignata, A, Landi, D, Navai, SA, et al. (2016). Tandem CAR T cells targeting HER2 and IL13Ralpha2 mitigate tumor antigen escape. The Journal of clinical investigation 126: 3036–3052.

35. Zah, E, Lin, MY, Silva-Benedict, A, Jensen, MC, and Chen, YY (2016). T Cells Expressing CD19/CD20 Bispecific Chimeric Antigen Receptors Prevent Antigen Escape by Malignant B Cells. Cancer Immunol Res 4: 498–508.

36. Asnani, M, Hayer, KE, Naqvi, AS, Zheng, S, Yang, SY, Oldridge, D, et al. (2019). Retention of CD19 intron 2 contributes to CART-19 resistance in leukemias with subclonal frameshift mutations in CD19. Leukemia.

37. De Giovanni, C, Nicoletti, G, Quaglino, E, Landuzzi, L, Palladini, A, Ianzano, ML, et al. (2014). Vaccines against human HER2 prevent mammary carcinoma in mice transgenic for human HER2. Breast Cancer Res 16: R10–R10.

38. Zgura, A, Galesa, L, Bratila, E, and Anghel, R (2018). Relationship between Tumor Infiltrating Lymphocytes and Progression in Breast Cancer. Maedica (Buchar) 13: 317–320.

39. Ruella, M, and June, CH (2018). Predicting Dangerous Rides in CAR T Cells: Bridging the Gap between Mice and Humans. Molecular Therapy 26: 1401–1403.

40. Blat, D, Zigmond, E, Alteber, Z, Waks, T, and Eshhar, Z (2014). Suppression of murine colitis and its associated cancer by carcinoembryonic antigen-specific regulatory T cells. Mol Ther 22: 1018–1028.

41. Baley, PA, Yoshida, K, Qian, W, Sehgal, I, and Thompson, TC (1995). Progression to androgen insensitivity in a novel in vitro mouse model for prostate cancer. J Steroid Biochem Mol Biol 52: 403–413.

42. Hull, GW, McCurdy, MA, Nasu, Y, Bangma, CH, Yang, G, Shimura, S, et al. (2000). Prostate Cancer Gene Therapy: Comparison of Adenovirus-mediated Expression of Interleukin 12 with Interleukin 12 plus B7-1 for *in Situ* Gene Therapy and Gene-modified, Cell-based Vaccines. Clinical Cancer Research 6: 4101.

43. Saffran, DC, Raitano, AB, Hubert, RS, Witte, ON, Reiter, RE, and Jakobovits, A (2001). Anti-PSCA mAbs inhibit tumor growth and metastasis formation and prolong the survival of mice bearing human prostate cancer xenografts. Proceedings of the National Academy of Sciences of the United States of America 98: 2658–2663.

44. Kern, P, Hussey, RE, Spoerl, R, Reinherz, EL, and Chang, H-C (1999). Expression, Purification, and Functional Analysis of Murine Ectodomain Fragments of CD8αα and CD8αβ Dimers. Journal of Biological Chemistry 274: 27237–27243.

45. Classon, BJ, Brown, MH, Garnett, D, Somoza, C, Barclay, AN, Willis, AC, et al. (1992). The hinge region of the CD8 alpha chain: structure, antigenicity, and utility in expression of immunoglobulin superfamily domains. Int Immunol 4: 215–225.

46. Kochenderfer, JN, Yu, Z, Frasheri, D, Restifo, NP, and Rosenberg, SA (2010). Adoptive transfer of syngeneic T cells transduced with a chimeric antigen receptor that recognizes murine CD19 can eradicate lymphoma and normal B cells. Blood 116: 3875–3886.

47. Park, AK, Fong, Y, Kim, SI, Yang, J, Murad, JP, Lu, J, et al. (2020). Effective combination immunotherapy using oncolytic viruses to deliver CAR targets to solid tumors. Sci Transl Med 12.

48. Kim, D, Pertea, G, Trapnell, C, Pimentel, H, Kelley, R, and Salzberg, SL (2013). TopHat2: accurate alignment of transcriptomes in the presence of insertions, deletions and gene fusions. Genome Biology 14: 1–13.

49. Anders, S, Pyl, PT, and Huber, W (2015). HTSeq—a Python framework to work with high-throughput sequencing data. Bioinformatics 31: 166–169.

50. Hsu, F, Kent, WJ, Clawson, H, Kuhn, RM, Diekhans, M, and Haussler, D (2006). The UCSC Known Genes. Bioinformatics 22: 1036–1046.

51. Mortazavi, A, Williams, BA, McCue, K, Schaeffer, L, and Wold, B (2008). Mapping and quantifying mammalian transcriptomes by RNA-Seq. Nat Meth 5: 621–628.

52. Lawrence, M, Huber, W, Pagès, H, Aboyoun, P, Carlson, M, Gentleman, R, et al. (2013). Software for Computing and Annotating Genomic Ranges. PLOS Computational Biology 9: e1003118.

53. Subramanian, A, Tamayo, P, Mootha, VK, Mukherjee, S, Ebert, BL, Gillette, MA, et al. (2005). Gene set enrichment analysis: A knowledge-based approach for interpreting genome-wide expression profiles. Proceedings of the National Academy of Sciences 102: 15545–15550.

54. Liberzon, A, Birger, C, Thorvaldsdóttir, H, Ghandi, M, Mesirov Jill P, and Tamayo, P (2015). The Molecular Signatures Database Hallmark Gene Set Collection. Cell Systems 1: 417–425.

55. Ashburner, M, Ball, CA, Blake, JA, Botstein, D, Butler, H, Cherry, JM, et al. (2000). Gene Ontology: tool for the unification of biology. Nat Genet 25: 25–29.

