## Supplemental Figures and Legends for "Pre-conditioning Modifies the Tumor Microenvironment to Enhance Solid Tumor CAR T Cell Efficacy and Endogenous Immunity"

**Supplemental Figures and Legends, Murad et al.**

Supplemental Figure 1

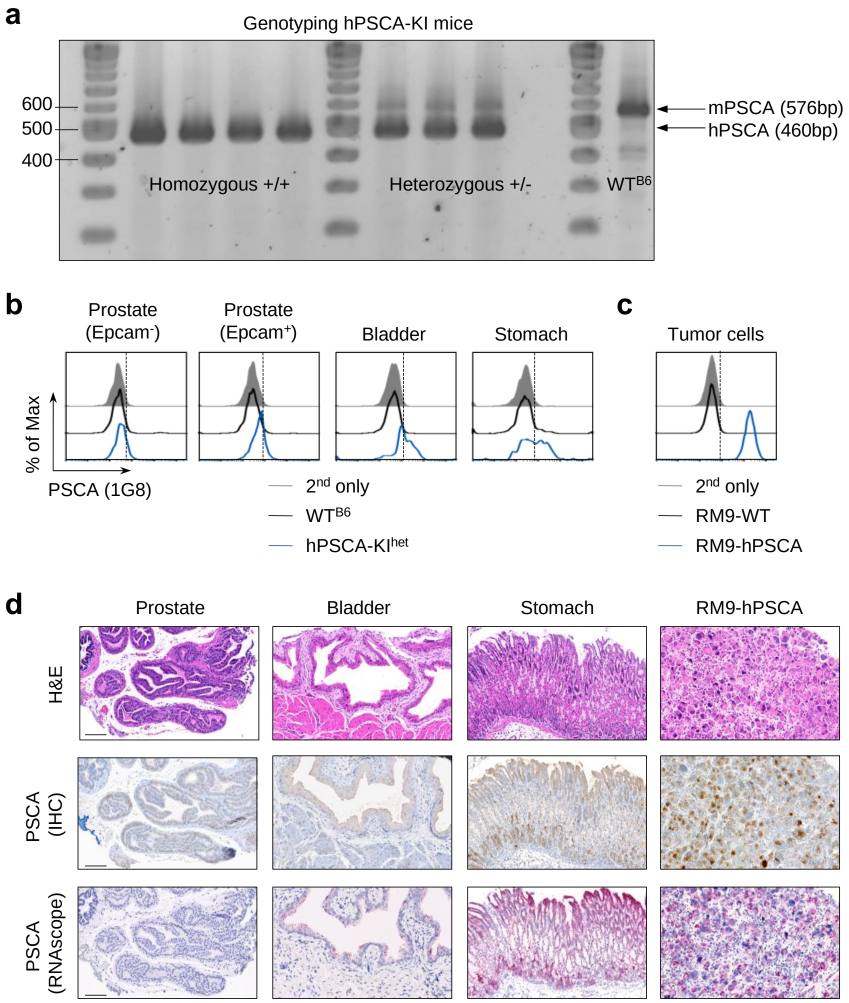


**Figure S1: hPSCA expression in normal tissues of hPSCA-KI mice.** a) Genotype analysis of hPSCA-KI mice (homozygous +/+, heterozygous +/-) for hPSCA knock-in as compared to wildtype C57BL6 ($WT^{B6}$) mice. Single bands at 576bp represents endogenous murine PSCA (mPSCA) and at 460bp the knock-in for human PSCA (hPSCA). b) Flow cytometry of hPSCA expression on single cell preparations of indicated normal tissues harvested from $WT^{B6}$ (black) and heterozygous (+/-) hPSCA-KI mice (blue) relative to controls. c) Flow cytometry of hPSCA expression in RM9-hPSCA tumor cells relative to controls. d) Immunohistochemistry (H&E and hPSCA) on normal prostate, bladder, stomach, and RM9-hPSCA tumors harvested from hPSCA-KI mice. Additional histological *in situ* hybridization analysis (RNAscope™) for hPSCA RNA expression (red dots) on normal tissues and RM9-hPSCA tumors. All images are at 20x magnification, scale bars represent 100 µm.

Supplemental Figure 2

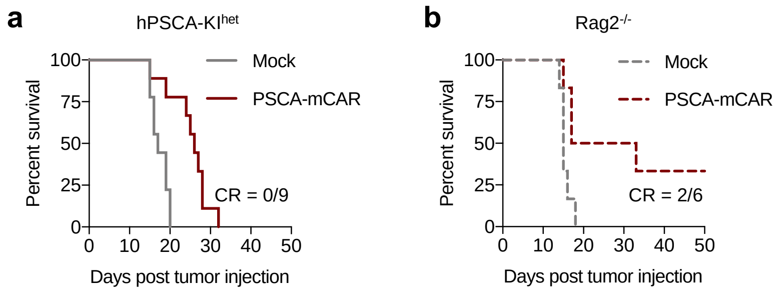


**Figure S2: PSCA-mCAR T cell therapy in RM9-hPSCA tumor-bearing immune competent hPSCA-KI and lymphoid deficient Rag2$^{-/-}$ mice.** Kaplan-Meier survival curves for RM9-hPSCA tumor bearing heterozygous PSCA-KI mice (a) and Rag2$^{-/-}$ (b) mice after intravenous (i.v.) administration of either $5.0x10^6$ Mock or PSCA-mCAR T cells. CR = complete curative response.

Supplemental Figure 3

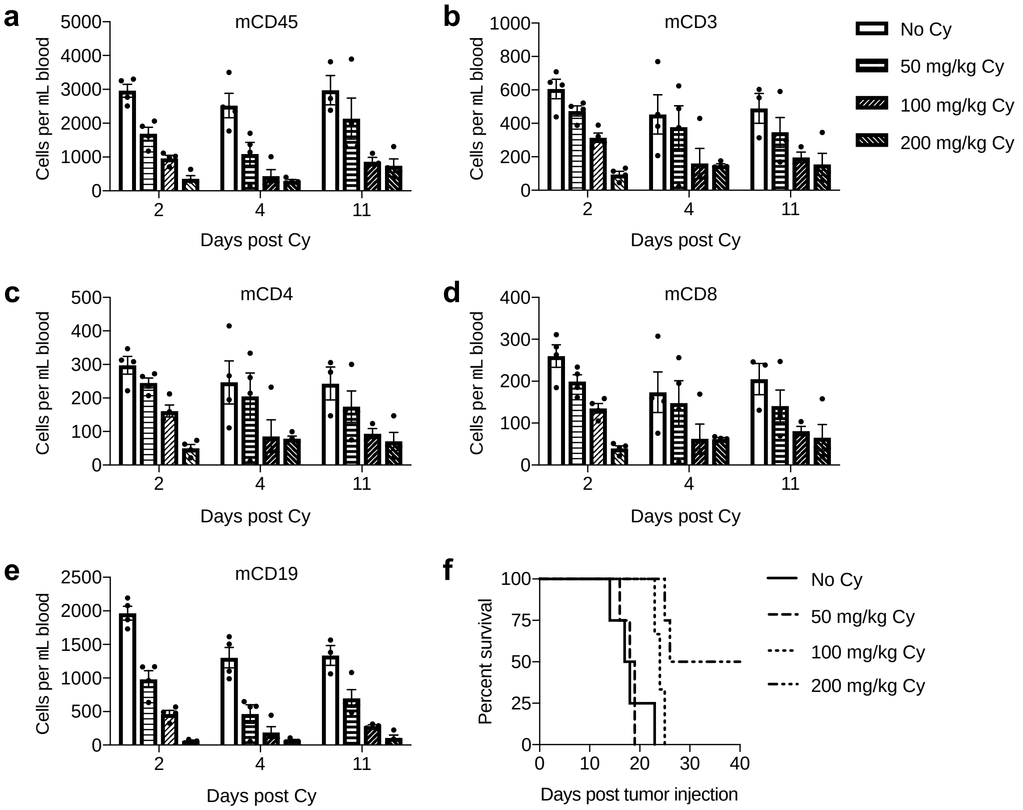


**Figure S3: Dose-dependent cyclophosphamide (Cy) depletion of immune populations and impact on overall survival of RM9-hPSCA tumor-bearing hPSCA-KI mice.** Flow cytometry of total murine CD45 (a), CD3 (b), CD4 (c), CD8 (d), and CD19 cells (e). Shown is absolute cells per µL of blood at days 2, 4 and 11 following a single i.p. administration of Cy at 0 mg/kg, 50 mg/kg, 100 mg/kg, or 200 mg/kg. f) Kaplan-Meier survival curve of hPSCA-KI mice engrafted intratibially (i.ti.) with RM9-hPSCA tumors and treated on day 3 post tumor injection with Cy (i.p.) at indicated doses.

Supplemental Figure 4

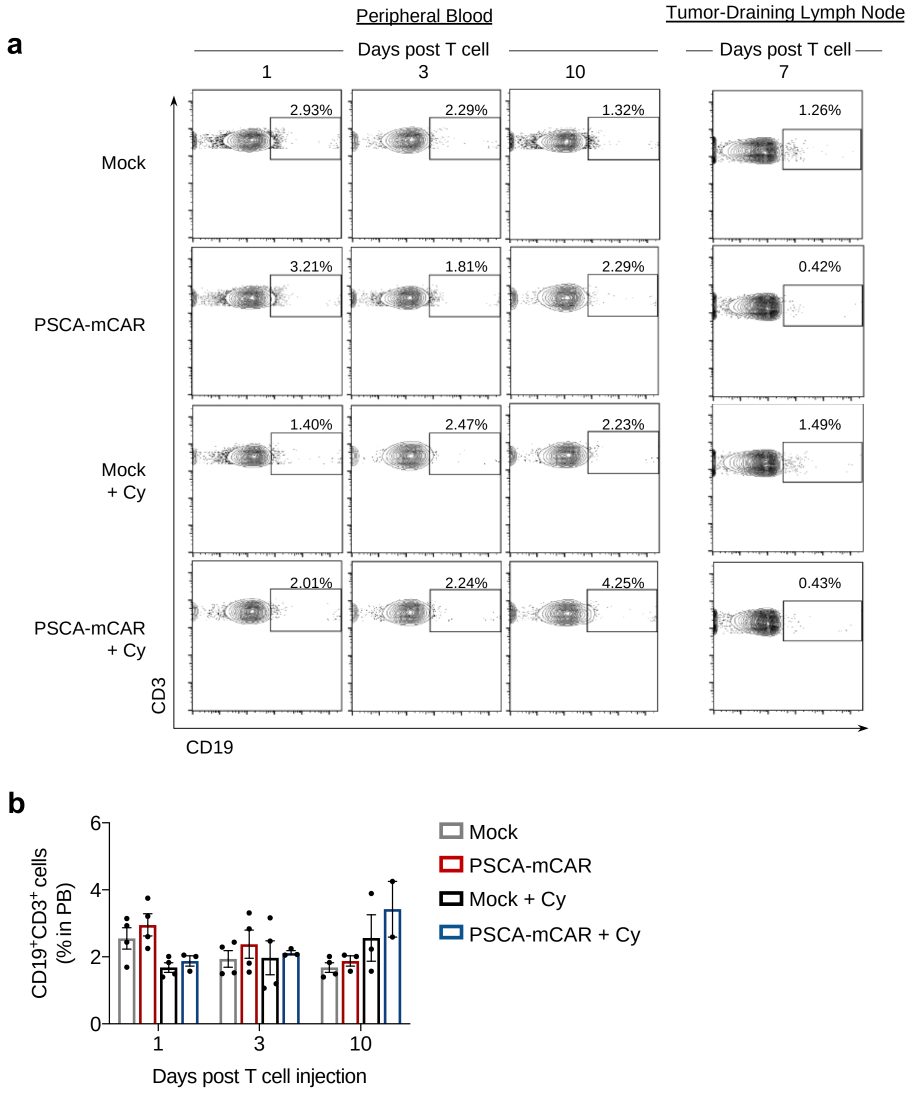


**Figure S4: Cy does not increase PSCA-mCAR T cell persistence in peripheral blood.** a) Representative flow cytometry of %CD19+CD3+ (%PSCA-mCAR, gated on CD45+CD3+) found in peripheral blood (PB) at days 1, 3 and 10 (left), and tumor-draining lymph nodes at day 7 (right), post T cell injection in RM9-hPSCA tumor bearing mice treated with Mock or PSCA-mCAR T cells with or without Cy pre-conditioning. b) Quantification of total %CD19+CD3+ (%PSCA-mCAR) found in PB of RM9-hPSCA tumor bearing mice treated with Mock or PSCA-mCAR T cells with or without Cy pre-conditioning at days 1, 3, and 10 post T cell injection.

Supplemental Figure 5

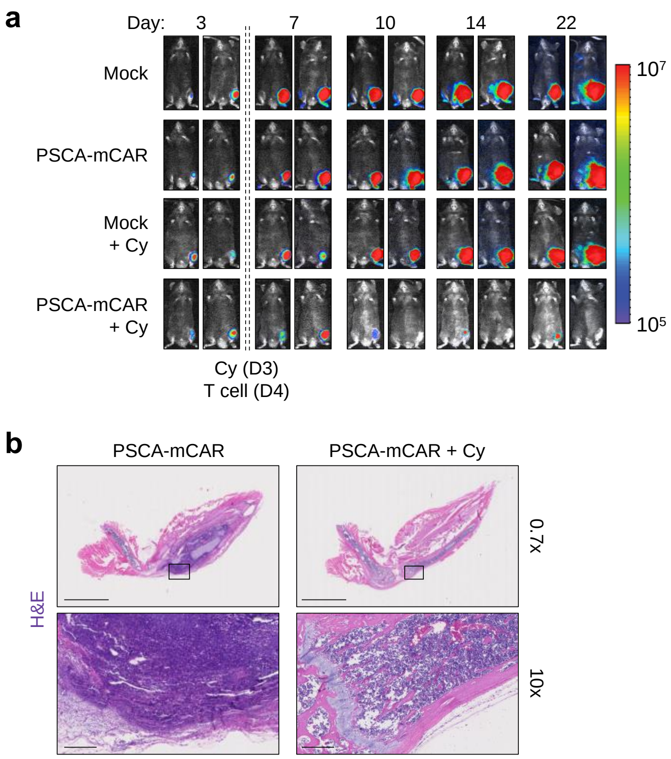


**Figure S5: Intratibial RM9-hPSCA tumor flux and histological analysis following Cy and PSCA-mCAR T cell treatment.** a) Representative tumor flux imaging for intratibial (i.ti.) engrafted hPSCA-KI mice showing two representative mice per treatment group at indicated days post tumor injection. b) Immunohistochemistry H&E staining of representative tibias harvested from RM9-hPSCA tumor bearing mice at day 7 post T cell injection highlighting efficacy of clearance of bone lesions in mice treated with PSCA-mCAR in combination with CPA pre-conditioning. 0.7x magnification scale bar represents 5 mm, 10x magnification scale bar represents 250 µm.

Supplemental Figure 6

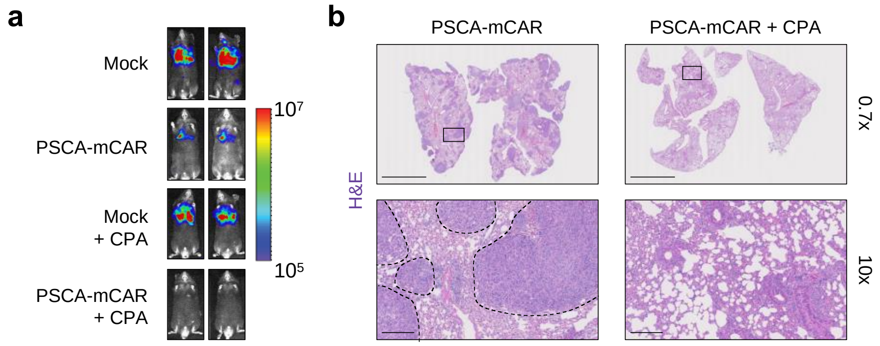


**Figure S6: Intrapulmonary KPC-hPSCA tumor flux and histological analysis following CPA and PSCA-mCAR T cell treatment.** a) Representative bioluminescent flux imaging of lung KPC-hPSCA tumors in hPSCA-KI mice at day 14 post T cell treatment with or without Cy pre-conditioning. b) H&E staining of representative lungs harvested from KPC-hPSCA tumor bearing mice at day 12 post T cell injection capturing early clearance of intrapulmonary KPC-hPSCA lesions (within dashed borders, 10x magnification) in mice treated as indicated. 0.7x scale bar magnification represents 5mm, 10x magnification scale bar represents 250µm.

Supplemental Figure 7

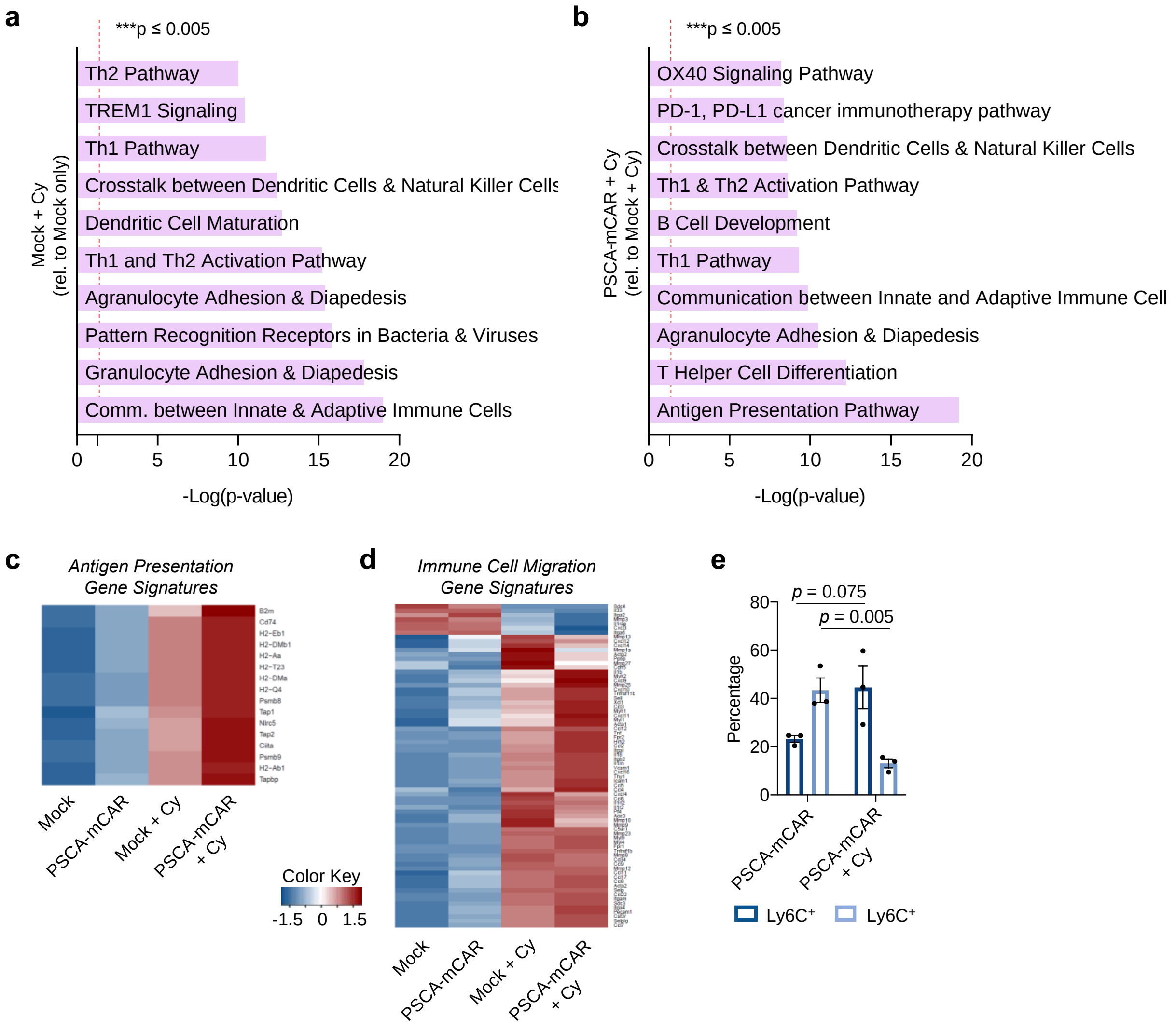


**Figure S7: RNA sequencing analysis of RM9-hPSCA tumors exhibit pro-immune transcript upregulation following Cy pre-conditioning and PSCA-mCAR T cell treatment.** Cy-mediated or PSCA-mCAR mediated impacts on immune related gene pathways as determined by Ingenuity Pathway Analysis (IPA). The top 10 significantly upregulated immune related gene pathways derived from bulk tumor RNA sequencing analysis shown as Mock + Cy treated mice relative to Mock T cell treatment alone (a) or PSCA-mCAR + Cy treated mice relative to Mock + Cy treatment alone (b). c ,d) Heatmaps showing standardized expression of RNA transcripts and gene signatures related to antigen presentation (c) or immune cell migration (d) in each indicated treatment group (scale -1.5 to +1.5). e) Flow cytometry quantification of frequency shift increase in peripheral blood monocytic pro-inflammatory Ly6C$^+$ (CD206$^-$F4/80$^+$) and decrease of granulocytic MDSC-like Ly6G$^+$ myeloid cells (gated on CD45$^+$CD11b$^+$) following PSCA-mCAR T cells with Cy pre-conditioning versus PSCA-mCAR T cells alone.
